# From Big Data to Small Scales: Machine Learning Enhances Microclimate Model Predictions

**DOI:** 10.64898/2025.12.01.691551

**Authors:** Alon Itzkovitch, Idan Sulami, Ronny Doron Efroni, Moni Shahar, Ofir Levy

**Author notes:** Corresponding author: Ofir Levy; mailing address: School of Zoology, Faculty of Life Sciences, Tel Aviv University, 6997801; ; fax: **;**.

## Abstract

1. Microclimates are critical for understanding how organisms interact with their environments, influencing behaviour, physiology, and species distributions. However, traditional physical heat-balance models for predicting ground temperatures in microhabitats often exhibit biases due to unaccounted environmental complexities and poorly constrained parameters. These limitations can hinder ecological research and conservation planning, particularly in the context of climate change.

2. In this study, we demonstrate how high-resolution drone-based mapping and machine learning can improve the accuracy of microclimate models. Using drone imagery, we generated detailed environmental maps, including solar radiation, vegetation indices, and skyview factors, to parameterize a physical heat-balance model. Validation with thermal maps derived from drone-mounted infrared cameras revealed systematic errors in the physical model’s predictions, including over- and underestimations under specific environmental conditions. To address these errors, we applied a random forest machine learning model to predict and correct biases in new prediction maps.

3. Our results show that machine learning reduced mean absolute errors by over 30% and mean square errors by 50%, while consistently narrowing the range of prediction inaccuracies. Key factors driving biases, such as vegetation cover, solar radiation, and height above ground, were identified, offering valuable insights for improving physical models. The machine learning corrections not only improved accuracy but also highlighted parameters and processes that were previously underrepresented or oversimplified in traditional models.

4. These findings illustrate the potential of combining machine learning with physical modelling to enhance microclimate predictions. This approach provides ecologists and conservation practitioners with a powerful tool to generate accurate, fine-scale microclimate maps, enabling better understanding of species responses to climate change and informing climate-resilient habitat management and conservation strategies.

## 1 INTRODUCTION

Climate conditions are fundamentally important in shaping how organisms experience their environment, from driving momentary impacts to influencing long-term evolutionary processes, and from guiding the decisions of individuals to structuring entire ecological communities (Potter et al., 2013). To understand the effects of climate on organisms, it is essential to examine the environmental conditions they experience in their immediate vicinity, often on scales ranging from meters to centimetres (Pincebourde and Woods, 2020). However, most available climate data are collected at much larger scales, often exceeding 10 km (Potter et al., 2013). Thus, to investigate the direct links between climate and the physiology, behaviour, distribution, and abundance of organisms, climate data with the fine spatial and temporal resolution are necessary (Kearney and Porter, 2017; Briscoe et al., 2023). To bridge the gap between large-scale climate data and the microclimates organisms experience, ecologists have developed microclimate models that translate regional climate conditions into localized microclimates (Kearney and Porter, 2017; Levy et al., 2016). These models, using both statistical and mechanistic approaches, aim to capture the heat balance between the ground, soil, and atmosphere near the ground at specific locations (Kearney and Porter, 2017; Levy et al., 2016; Zanchi et al., 2023; Mukonza and Chiang, 2022). However, parameterizing, validating, and especially reducing the biases of these models across wide, high-resolution spatial scales remain challenging (Briscoe et al., 2023).

Recent technological and algorithmic advancements are now helping to address these challenges. On one hand, the availability of off-the-shelf sensors, such as thermal and standard cameras mounted on drones, has enabled cost-effective and extensive data collection, providing large datasets for both microclimate model input and output validation. On the other hand, this wealth of data can be leveraged to train Machine Learning (ML) models, that can predict climate or reduce biases in climate model predictions, including temperature, precipitation, and extreme weather events (reviewed by Levy and Shahar, 2024). In particular, with sufficient data, ML models can learn complex relationships between model outputs, input, and observed data, thereby enhancing the reliability of predictions—a critical factor for ecosystem management, agricultural planning, and urban development (Chen et al., 2023; Sabarinath et al., 2023; Kesavavarthini et al., 2023; Zhu et al., 2022; Han et al., 2024).

Machine Learning models can serve for two purposes: either directly predict climate conditions or correct the bias of physical-based climate models (Levy and Shahar, 2024). In direct prediction, ML models estimate microclimatic conditions solely from available environmental inputs, potentially replacing the physical model. However, it may be challenging for such models to extrapolate under unfamiliar conditions, an important caveat when predicting the future climates (Hernanz et al., 2024). In bias correction, ML is used to learn and adjust for systematic errors in the output of a physical model, effectively combining the strengths of process-based modelling with data-driven correction (reviewed by Levy and Shahar, 2024). Recent ML models have significantly reduced biases in surface temperature and precipitation forecasts generated by global physical models (Sabarinath et al., 2023; Kesavavarthini et al., 2023). These models often use landscape characteristics, such as elevation, as features and correct biases in the physical models’ predictions (Levy and Shahar, 2024). Despite the demonstrated success of ML for large-scale climate models, bias correction has not yet been widely applied to microclimate models. However, recent studies demonstrate that ML can accurately predict microclimatic variables—including temperature and relative humidity (Han et al., 2024; Zanchi et al., 2023; Arulmozhi et al., 2021), suggesting that ML models can effectively capture micro, non-linear spatial and temporal patterns across a wide range of environmental contexts (Han et al., 2024; Zanchi et al., 2023; Mukonza and Chiang, 2022). Therefore, ML models may also have a strong potential to not only predict microclimates, but also learn to correct biases in the physical models’ prediction.

In this study, we used drone-based data collection to develop a physical heat-balance microclimate model. We then applied a machine-learning model to correct its prediction biases (systematic errors, such as consistent over- or underprediction under specific conditions). We validated both physical and machine-learning models using thermal maps derived from drone surveys conducted across different seasons and times of day in a desert habitat. We compared the performance of the models and analyzed the importance of various variables for bias correction, identifying parameters or physical processes that may be inaccurately or incompletely represented in the microclimate model. We hypothesized that machine learning could reduce errors in microclimate estimates, as well as diminish the relationships between input variables and model errors.

## 2 MATERIALS AND METHODS

### 2.1 General Approach

Our modelling approach combines a physically-based microclimate model with a machine learning (ML) bias-correction model. The physical model explicitly simulates the ground surface energy balance—including solar and longwave radiation, sensible and latent heat fluxes—to predict ground temperatures at high spatial resolution. The ML model is then used to estimate and correct systematic biases in the physical model’s predictions, thereby improving overall accuracy.

We parameterized the models by collecting drone imagery through mapping missions conducted across various seasons and times of day. Our drone was equipped with a dual-sensor camera capable of capturing both RGB (visible light) and infrared (IR) images. We used these images to generate RGB and Digital Surface Models (DSMs) maps at a resolution of 3 cm^2^, and thermal maps at a resolution of 15 cm^2^. For each flight, we created input maps for the micro-climate model, including maps of shade conditions and solar radiation, using the DSM and meteorological data. We calculated ground temperatures at a fine spatial resolution of 15 cm^2^ per pixel by feeding these input maps and meteorological data—obtained from both a mobile meteorological station and an online database—into the microclimate model. Finally, we trained a machine learning (ML) model to correct errors in our microclimate predictions. This was done by comparing the ground temperature predictions from the microclimate model to the actual temperatures recorded on the drone-based thermal maps. The ML model learned to estimate the difference (or error) between predicted and observed temperatures, enabling the reduction of bias and improving the accuracy of microclimate predictions. A visual description of the data collection, microclimate modelling, and training of the ML correction model is shown in Fig. 1.

**Figure 1:**
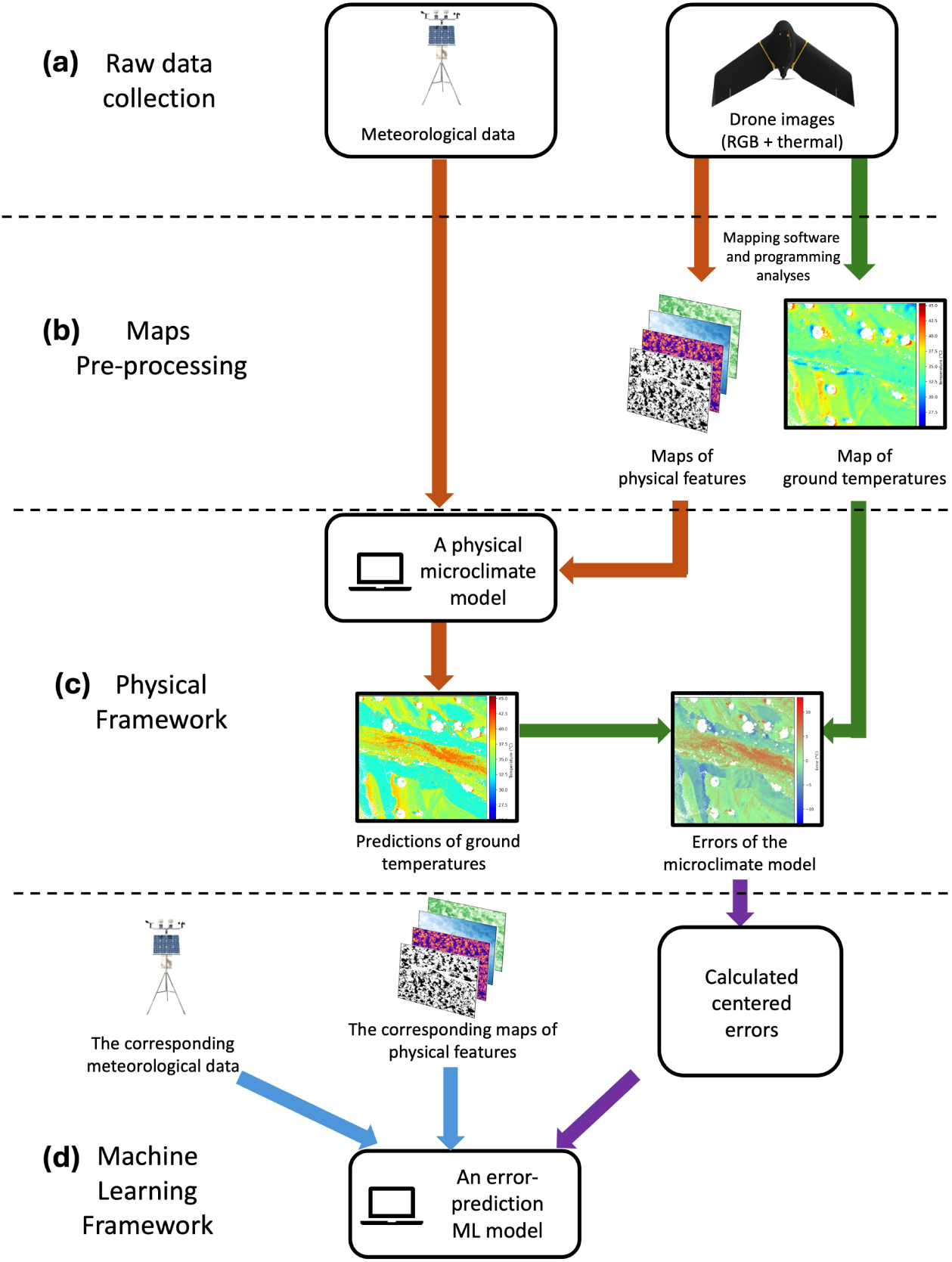
Workflow of the parameterization of the physical microclimate model and the training of the ML bias correction model. (a) Data are collected using drones and meteorological stations or online databases. (b) Drone images are processed into high-resolution physical feature maps (e.g., shade and solar radiation) and a ground temperature map for bias correction. (c) The physical microclimate model uses physical feature maps and meteorological data to predict ground temperatures (orange arrows). The predictions are compared with real ground temperatures to generate an error map (green arrows). (d) A machine learning model is trained on centralised error maps (purple arrows) to predict model errors for new data (blue arrows).

### 2.2 Drone-based Microclimate Model

We employed a physical microclimate model to estimate ground temperature using surface characteristics and meteorological variables, including air temperature, pressure, and albedo. Ground temperature calculations for each pixel were based on physical heat-balance equations derived from the NOAH Microphysics model (Niu et al., 2011), following the approach described by Levy et al. (2016). A detailed description of the model is in the online supplementary.

### 2.3 Model Parameterization

We parameterized the model using meteorological variables, including soil, ground, and air temperatures, as detailed in Table S1. These parameters were either measured directly in the field or obtained from the Global Land Data Assimilation System (GLDAS, Rodell et al., 2004). Parameters were assumed to be constant within each map, except for solar radiation, shade, skyview, greenness, and height above bare ground, which varied between pixels (see below). Importantly, the measured ground temperature from the meteorological station served exclusively as the initial condition for each pixel-specific microclimate simulation.

To capture these spatial variations accurately, we collected spatial data through drone mapping flight missions and simultaneous meteorological data collection. The drone mapping aimed to characterize differences between pixels in terms of solar radiation, shade, skyview, greenness, and height. Meanwhile, the meteorological data provided coarse-resolution climate information typically used to model heat load differences across the habitat. This integrated approach allowed us to account for both fine-scale spatial heterogeneity and larger-scale climate drivers. From June 2019 to May 2021, we conducted 33 drone flights over 8 days, spanning morning to evening hours. A comprehensive list of flight dates and times is provided in Table S2.

#### 2.3.1 Field surveys

##### Study area

We collected field data in the Judean Desert in Israel (31°28’N, 35°10’E), bounded to the east by the Dead Sea, 400 m below sea level. The region is classified as a hot desert (BWh) according to the Köppen-Geiger system and receives less than 100 mm of annual rainfall. Flora is predominantly perennial shrubs and annual grasses, which vary in density. Our study site was in Parking Tse’elim River (31°21’04.8”N 35°21’11”E), a rocky habitat with very sparse vegetation. Temperatures in the Judean Desert vary seasonally, and microclimates play a major role in the ecology and physiology of animals. During summer, mean ground temperatures in the open range from 30°C in the early morning to 44°C at noon; shade cover offers substantial thermal shelter—maximal ground temperatures reach only 37 °C under rocks. Winter temperatures are cooler, with ground temperatures reaching 27°C in the open and 25°C in the shade. All surveys were conducted in a natural desert habitat during daytime hours due to logistical constraints and local restrictions on drone operation at night.

##### Meteorological data

At the beginning of each field day, we placed a mobile meteorological station (MaxiMet GMX501 Compact Weather Station, GILL instruments, United Kingdom) in a sun-exposed and horizontal location (i.e., <10 degrees slope) within our mapping area. Recorded meteorological conditions included solar radiation, air temperature at 1.2 m height, ground temperature, wind speed, wind direction, relative humidity, and air pressure at 10-minute intervals.

##### Online meteorological data

We used the Global Land Data Assimilation System (GLDAS, Rodell et al., 2004) dataset to extract variables that are not available from the meteorological station but are needed for parameterizing the microclimate models, such as albedo and soil moisture (see Table S1).

##### Drone Data

Our framework uses drone images to generate input and validation (thermal) maps for the microclimate model. We employed an eBee X mapping drone (SenseFly, Switzerland) equipped with a dual RGB and thermal camera (Duet T, SenseFly, Switzerland, with an accuracy of ±5°C). To ensure our data covered a variety of conditions, we conducted surveys at different times of day and throughout various seasons (Table S2). Each mapping session was planned using Flight Planning Software (eMotion 3, SenseFly, Switzerland) and covered areas ranging from approximately 0.3 km^2^ to 1 km^2^, with 65% overlap between images.

During each session, the software facilitated the manual launch of the drone, navigated the drone through the survey area, captured images, and returned the drone to a designated landing point. Each flight lasted between 20 to 60 minutes. The images were stored on an SD card and transferred to a laptop after each flight. Notably, this approach is accessible even to owners of standard, off-the-shelf drones equipped with RGB and/or thermal cameras. Mapping can be easily planned and executed using free flight-planning apps, such as Pix4DCapture (Pix4D Inc., Denver, CO, USA), making microclimate modelling feasible without requiring a high-end drone like ours.

#### 2.3.2 Map Development

After collecting the drone images and meteorological data, we developed the necessary maps for the microclimate models in two main stages. In the first stage, drone images were processed using professional photogrammetry software (Pix4DMapper version 4.23, Pix4D Inc., Denver, CO, USA) to generate high resolution maps. From the RGB images, the software produced Digital Surface Models (DSMs, m), Digital Terrain Models (DTMs, m), and RGB orthophotometric maps at a resolution of 3 cm (pixel size = 3 cm^2^). These maps can also be created using the freely available Open Drone Map software (Gbagir et al., 2023). The thermal images were used to generate a thermal map at a 15 cm resolution (pixel size = 15 cm^2^), which was employed for validating model predictions and training the bias-correction model. Since the thermal camera has an accuracy of ±5°C, we calibrated each map based on our meteorological station. Specifically, for each pixel, we subtracted the calibration value, which was the difference between the temperature recorded at the meteorological station and the temperature at the corresponding coordinates on the thermal map, assuming a uniform error across the map.

We resampled all maps from each flight to a 15 cm resolution to match the thermal map. Moreover, to balance data quantity and computational cost, we then randomly selected five maps per flight for further analysis. The size of each sampled map was 1024×1024 pixels, which is approximatly 150 m^2^). Despite the drone GPS having an accuracy of approximately 2 m, maps generated from images of the same flight were perfectly aligned.

In the second stage, DSM, DTM, and RGB maps were further analyzed to create input rasters for the microclimate model, including solar radiation (W m^–2^), shade (yes/no), skyview (%), height (m), and vegetation cover (Triangular Greenness Index, TGI; Starý et al., 2020). These maps represent environmental conditions at a single point in time—specifically, the moment of the drone flight—capturing solar radiation and shade as they occurred during the survey. Notably, our shade map did not include areas directly under vegetation, as below-canopy pixels are not visible to the drone’s thermal camera. Thus, in our analysis, shaded pixels refer exclusively to bare ground (TGI<0.04) areas that, at the time of each flight, were shaded by nearby objects—such as rocks, sparse vegetation, or other landscape features—due to the sun’s position during the survey.

Raster maps were generated using a combination of R scripts and GRASS GIS commands (GRASS Development Team, 2022, ver. 7.4.0) called from within R (R Core Team, 2021, ver. 4.1.0), utilizing the *rgrass7* package (Bivand, 2000). Both stages are shown in Fig. S1. A detailed description of the maps’ creation is in the online supplementary.

### 2.4 Machine Learning Bias-correction Model

All models, including microclimate models, exhibit some degree of bias. Machine learning (ML) approaches are increasingly used to correct such biases in climate model predictions, including temperature, precipitation, soil moisture, and extreme events (Chen et al., 2023; Sabarinath et al., 2023; Kesavavarthini et al., 2023; Zhu et al., 2022; Kratzert et al., 2019). In this study, we implemented a random forest regression model using the scikit-learn package in Python (Pedregosa et al., 2011) to reduce bias in our microclimate predictions.

We selected random forest because of its proven effectiveness in capturing non-linear relationships and complex interactions among predictors (Thessen, 2016), as well as its robustness and widespread adoption. While the random forest does not explicitly represent physical energy balance mechanisms, it serves as a bias-correction layer on top of the physically-based model. By leveraging empirical relationships between environmental variables and model prediction errors, it helps to reduce systematic bias in microclimate predictions. This hybrid approach preserves the mechanistic foundation of the physical model while harnessing the predictive power of machine learning.

To train the ML model, we first calculated the error for each pixel as the difference between the microclimate model’s prediction and the observed temperature. These errors served as the target variable for the random forest, which was trained using the same input maps as the microclimate model (shade, solar radiation, skyview, and height above ground), along with the raw predictions from the physical model, since model errors may vary with temperature. To avoid the ML model simply learning systematic errors from the physical model, we used a centered error for each pixel (the microclimate model error minus the mean prediction error across the map, denoted as *mPE*) as the target variable.

The random forest was configured with 100 estimators, a mean squared error loss function, a maximum tree depth of 10, the default minimum samples required to split an internal node, and bootstrap sampling enabled. To rigorously evaluate model performance and prevent information leakage, we divided our dataset by day, with 80% of the data (115 maps from 23 flight surveys) used for training and 20% (50 maps from 10 additional surveys) for testing. We assessed the ML model by predicting the error for each pixel in the test maps and comparing the original error of the microclimate model to the error after applying the ML-predicted correction.

To compare biases and overall prediction errors with and without ML correction, we calculated the mean error (ME), mean squared error (MSE), and mean absolute error (MAE) for the physical model’s predictions (*e_micro_*) and after the ML correction, both with and without incorporating the mean prediction error (*mPE*) (See flowchart of model usage and validation in Fig. 2). These metrics were calculated for each map as follows:

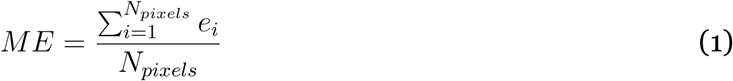

**Figure 2:**
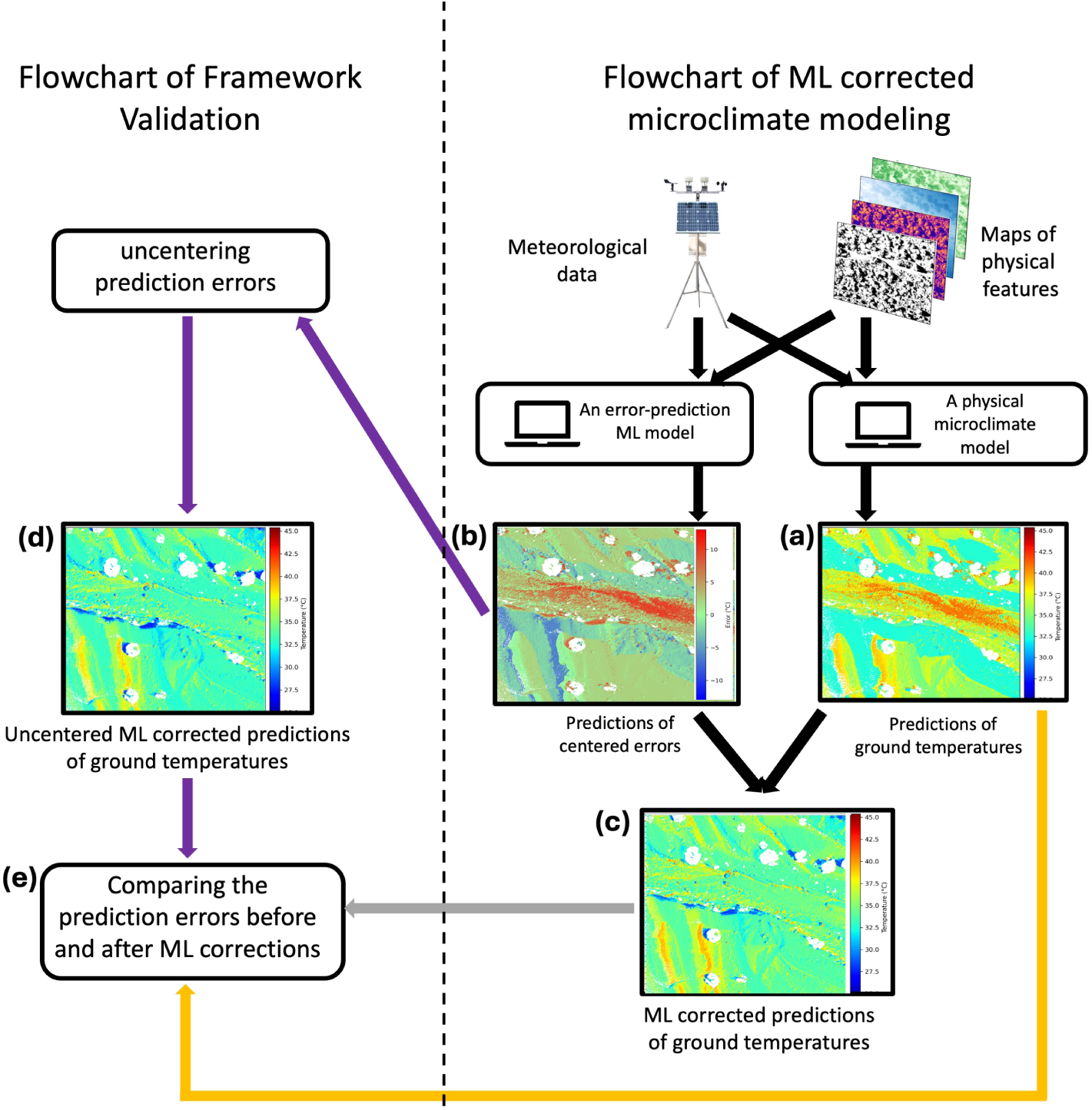
Flowchart of the ML correction usage (right) and our validation process (left). To use the ML correction model (see Figure 1 for training flowchart), meteorological data and physical feature maps (e.g., shade and solar radiation) are used to parameterize the physical microclimate model to calculate the ground temperature at each pixel (a) and by the ML correction model to predict the centred prediction error at each pixel (b). The combination between the two maps is the ML-corrected predictions (c). In our validation, we also calculated the uncentralized ML- corrected predictions (e). Our final validation analysis included both original microclimate predictions (orange arrow) and the corrected predictions using both centralised (grey arrow) and un-centralised (purple error) ML corrections errors (purple arrows). These three maps were then compared to the observed thermal map obtained from the drone (e).

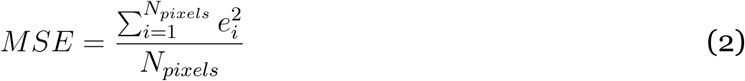

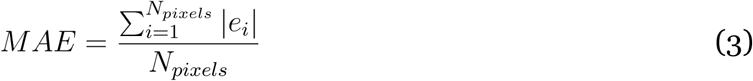

Here, *e_i_* represents the model error, which could be the error before applying the machine learning (ML) correction, or the error after applying the ML correction, with or without incorporating the mean prediction error (*mPE*).

Since pixels’ temperatures under canopy cover were not visible to the drone, we excluded such pixels from our analysis (pixels with TGI>0.04). Thus, validation of shaded pixels refers to bare ground pixels that were temporarily shaded by nearby objects, rather than being covered directly by vegetation.

### 2.5 Statistical Analysis

We used linear regression models to test our hypotheses, employing Bayesian analysis to evaluate the effectiveness of the ML correction model in reducing model errors and to assess the influence of various variables on prediction errors, both with and without ML correction.

First, we examined the effect of our ML correction model on mean error (ME), mean absolute error (MAE), and mean square error (MSE), both before and after ML correction, and with or without incorporating *mPE* after correction. We fitted a Linear Mixed Effects Model for ME and Generalized Linear Mixed Effects Models for MAE and MSE, using a Gamma distribution with a log link function for the latter two. To account for repeated measurements from the same maps, we included map names as a random effect in the models.

Second, we analyzed how the model features (i.e., explanatory variables) influenced possible systematic biases (e.g., overpredicting with increased solar radiation) of the microclimate model, both with and without ML correction. We selected 100 pixels from each testing map, ensuring at least 15 meters of separation between pixels to reduce spatial correlation. For these pixels, we fitted two Linear Mixed Effects Models with solar radiation, TGI, skyview, and height as continuous explanatory variables, shade as a categorical factor, and microclimate model error as the response variable, either before or after ML correction. For simplicity, we included only the interactions between the continuous variables and shade, as shade strongly affects microclimate. Map names were included as random effects to account for repeated measures, and we incorporated a variance structure to handle heterogeneity in the data. We determined the optimal random and variance structures using the *lme* function from the *nlme* R package (Pinheiro and Bates, 2000) by first comparing models with different random structures and then different variance structures, following the guidelines in Zuur et al. (2009). Models were compared using Akaike Information Criterion (AIC) values (Burnham and Anderson, 2002).

We fitted the models using the PyMC Python package (Salvatier et al., 2016) and ran them with a No-U-Turn (NUTS) sampler (Homan and Gelman, 2014) with over 1000 iterations and 4 chains. Convergence was assessed through trace plots, the Gelman-Rubin *R*^^^diagnostic, and effective sample size. To evaluate the significance of predictors, we examined the posterior distributions of the coefficients and calculated their 95% credible intervals. A coefficient was considered significantly different from zero at the 5% level if its high-density interval (HDI) did not include zero.

For regression models with log link functions, parameter estimates are presented as percentages, representing the relative change in microclimate model errors. These percentages were calculated as 100 *· e^k^*, where *k* is the estimated parameter (Ntzoufras, 2009).

## 3 RESULTS

### 3.1 Comparison of Microclimate Biases Before and After Machine Learning Correction

Our drone-based physical microclimate model performed well, and the machine learning correction significantly reduced bias in the model’s predictions.

The mean error (ME) of the physical model predictions was 0.56 *±* 1.97°C (mean *±* SD; Table S3). According to our linear model, applying the machine learning correction significantly reduced the mean error, decreasing it by 0.911 *±* 0.332°C, resulting in a tendency for the model predictions to slightly underestimate temperatures. When we added back the mean prediction error (*mPE*)—the average error across all pixels—to the machine learning-corrected predictions, the mean error was reduced by 1.471 *±* 0.334°C, with no significant difference between the two correction methods (Table 1A). Additionally, the machine learning correction alone improved error values in 62% of the test maps, while the correction with mPE added back showed improvement in 64% of the test maps (Table S3).

**Table 1:** Statistical results for the mean error (ME), mean absolute error (MAE), and mean squared error (MSE). We show the estimated mean error of the physical model (Intercept) and the effect of applying machine learning correction, with and without adding back the mean prediction error (mPE). For each metric, the table reports the mean *±* SD and 95% confidence intervals (CI 2.5% and 97.5%).

| **Metric** | **Mean ± SD** | **CI 2.5%** | **CI 97.5%** |
| --- | --- | --- | --- |
| **(A) ME** |  |  |  |
| Intercept | 0.551 ± 0.282 | 0.008 | 1.117 |
| Without mPE | -0.911 ± 0.332 | -1.583 | -0.251 |
| With mPE | -1.471 ± 0.334 | -2.100 | -0.791 |
| Difference Between ML Corrections | 0.560 ± 0.286 | -0.017 | 1.109 |
| **(B) MAE** |  |  |  |
| Intercept | 0.995 ± 0.040 | 0.918 | 1.073 |
| Without mPE | -0.379 ± 0.056 | -0.491 | -0.269 |
| With mPE | -0.413 ± 0.057 | -0.523 | -0.302 |
| Difference Between ML Corrections | -0.023 ± 0.042 | -0.105 | 0.059 |
| **(C) MSE** |  |  |  |
| Intercept | 2.435 ± 0.062 | 2.314 | 2.555 |
| Without mPE | -0.750 ± 0.114 | -0.971 | -0.526 |
| With mPE | -0.739 ± 0.098 | -0.927 | -0.543 |
| Difference Between ML Corrections | 0.005 ± 0.046 | -0.089 | 0.093 |

The mean absolute error (MAE) and mean square error (MSE) of the physical model predictions were 2.75 *±* 0.849°C and 12.129 *±* 6.612°C^2^, respectively (Table S3). For MAE, our Gamma linear model indicated that applying the machine learning correction reduced the error by 31.4 *±* 3.9%, and by 33.7 *±* 3.8% when mPE was added back (Table 1B). These corrections improved the MAE in 80% and 84% of the test maps, respectively (Table S3B). For MSE, the machine learning correction reduced the error by 52.5 *±* 5.5%, and by 52.0 *±* 4.8% when mPE was added back (Table 1C), with improvements observed in 82% and 88% of the test maps, respectively (Table S3).

Finally, we found that the machine learning correction centralized and narrowed the distribution of model biases (see Fig. 4). Before applying the machine learning correction, 90% of the errors in the physical model predictions were within *±* 6°C, 95% were within *±* 7°C, and 99% were within *±* 9°C. After the machine learning correction, 90% of the corrected errors were within *±* 4°C, 95% were within *±* 5°C, and 99% were within *±* 7°C. A similar effect was observed when the mean physical error was added back to the machine learning-corrected predictions (Fig. 4).

In summary, both machine learning corrections significantly reduced the errors of the physical model across all metrics (mean error, mean absolute error, and mean squared error), and the improvements were consistent across the majority of test maps. The effect of adding back *mPE* was not significant, indicating that adding *mPE* did not provide an additional benefit over correcting the physical model errors alone.

### 3.2 The Effect of ML Correction on Systematic Biases

We found that model features significantly affected the biases in the physical model, and the ML correction substantially reduced the magnitude of several of these effects (Fig. 5). Importantly, these relationships differed between the open and shaded microhabitats. For the physical model (Table S4A), in the open microhabitat, the model overestimated temperatures under high solar radiation and skyview conditions, while underestimating temperatures in areas with higher vegetation cover (TGI) and at greater heights. In contrast, in the shaded microhabitat, the overestimation due to solar radiation was more pronounced, and the effect of skyview reversed, leading to underestimation in areas with high skyview. Neither TGI nor height significantly affected errors in the shaded microhabitat.

**Figure 3:**
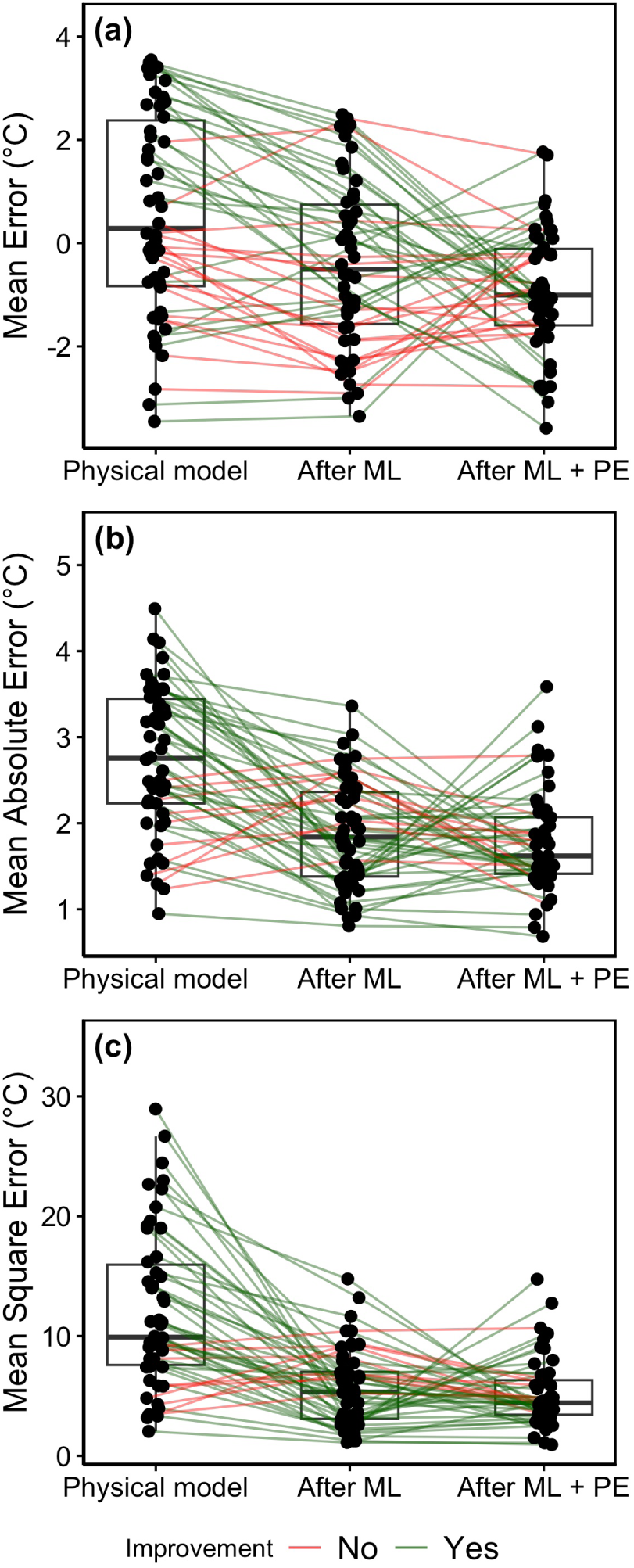
Machine learning decreased model errors. We show the distribution of the model mean errors (a), mean absolute error (b), and mean square error (c) of the original physical model and after our ML correction. Each dot represents a map, with lines connecting data of the same map. The green and red lines represent maps in which the ML correction improved or worsened the physical model predictions, respectively.

**Figure 4:**
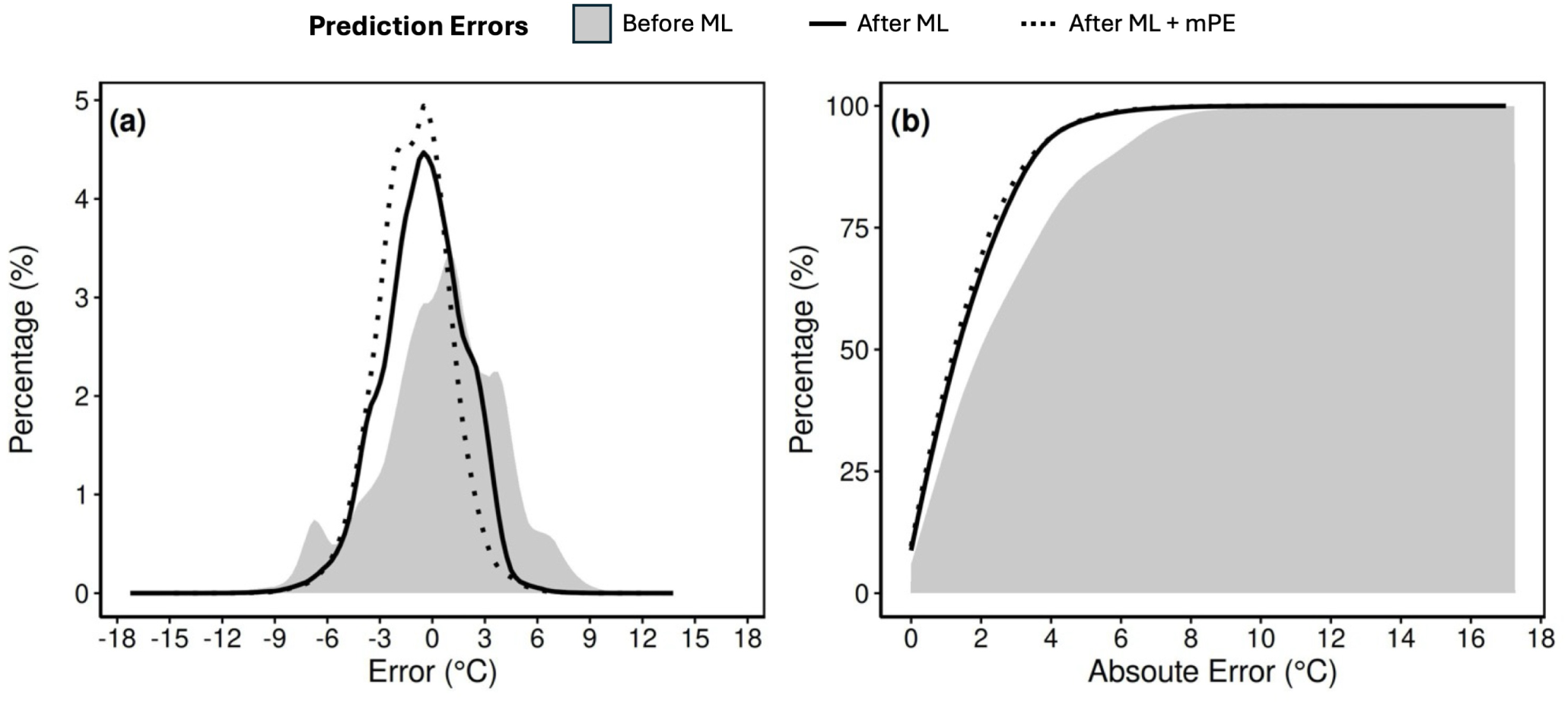
After the machine learning corrections, the microclimate errors are closer to zero. We show both (a) regular and (B) cumulative histograms of errors (a - errors, b - absolute errors) before and after our ML correction.

**Figure 5:**
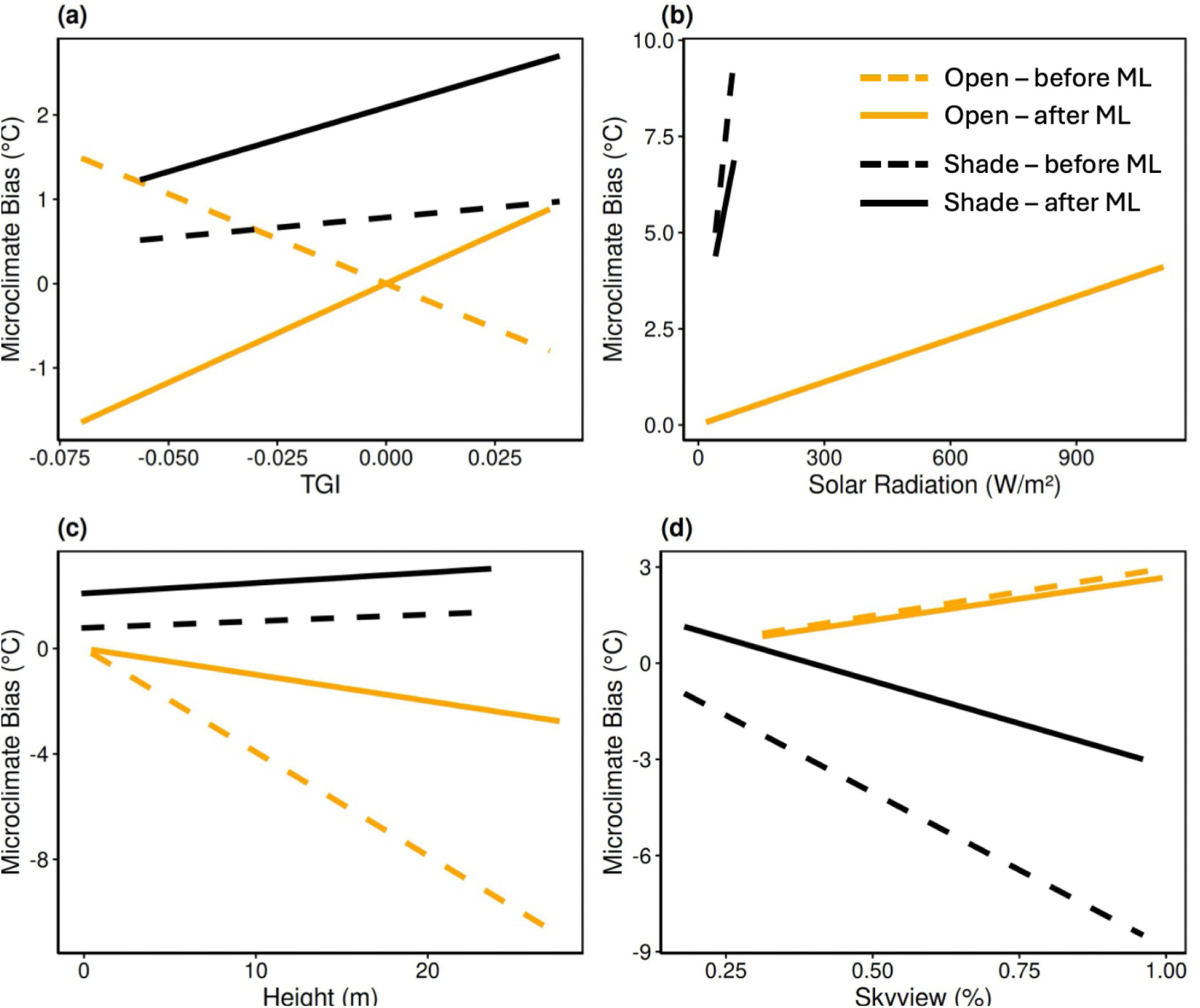
The effect of the machine learning (ML) correction model on systematic biases in microclimate predictions. Each panel presents the fitted regression slope between microclimate bias (°C) and a given climatic parameter, calculated from our statistical analysis, before (dashed lines) and after ML correction (solid lines). Parameters shown are: (a) Triangular Greenness Index (TGI), (b) Solar Radiation (W/m²), (c) Height (m), and (d) Skyview (%). Slopes are derived using 100 pixels from each test map. ML correction reduces the magnitude of these slopes, indicating that bias becomes less sensitive to environmental variation.

The ML correction significantly altered the relationships between model features and errors (Fig. 5, Table S4B). Notably, the correction reversed the effect of TGI, which shifted from causing underestimation to overestimation at higher TGI values in both the open and shaded microhabitats. In the shaded microhabitat, the ML correction reduced the effect of skyview, but TGI and height remained non-significant. However, solar radiation retained a significant positive effect on model errors after the correction.

When comparing the magnitude of slopes before and after the ML correction (Table S4C), the correction substantially improved several features. For example, the slope for height in the open microhabitat was significantly reduced, indicating a reduction in bias. In the shaded microhabitat, the correction improved the model’s performance with respect to solar radiation and skyview, yielding more accurate estimates.

Additionally, visual inspection revealed that the ML corrections successfully reduced over-estimation and underestimation of predicted temperatures (see red pixels in Fig. 6A, and B, and Figs. S6–S8).

**Figure 6:**
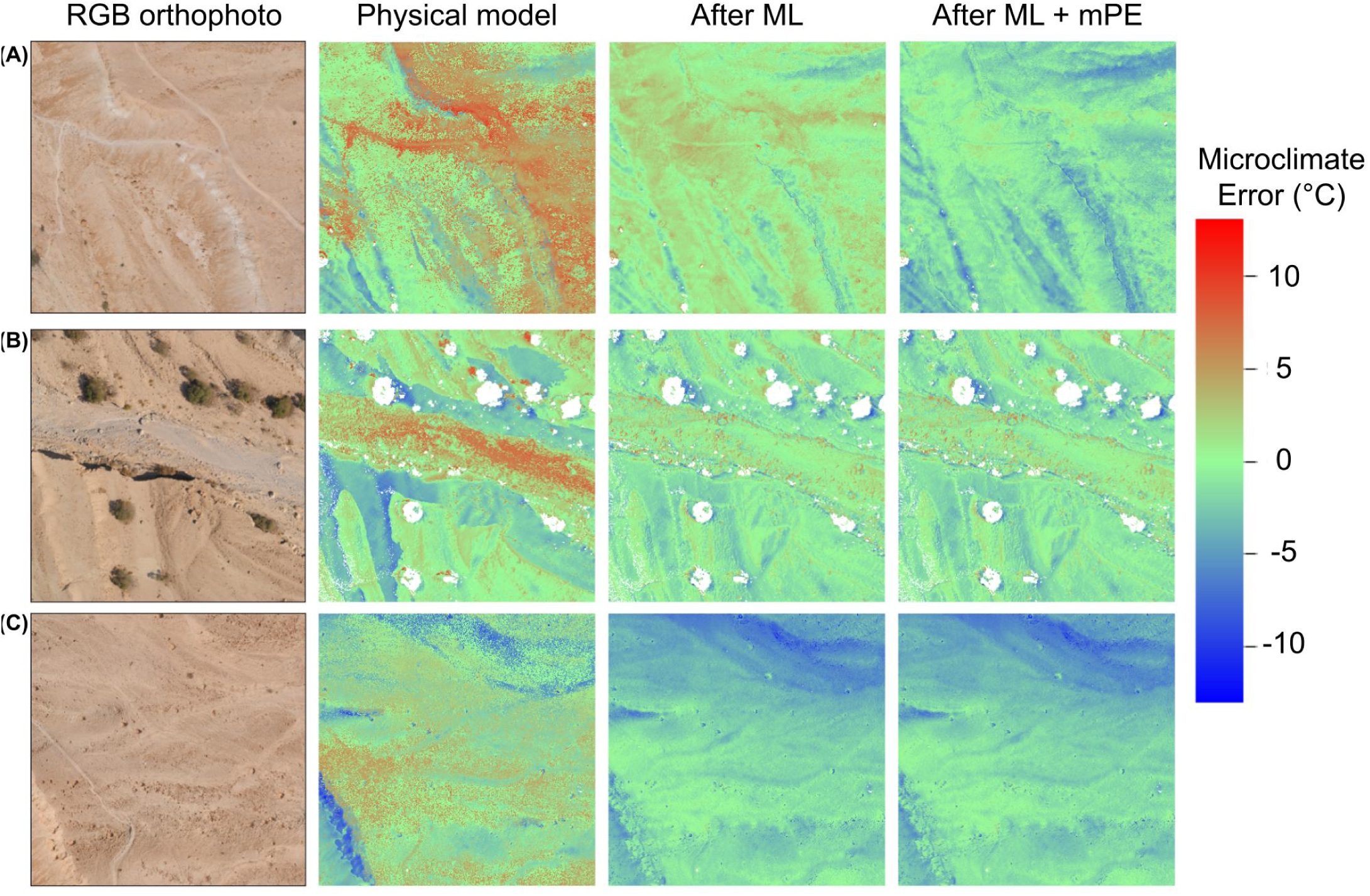
Sample maps showing the effect of ML correction on the bias of the physical model. Each row represents one map, and the columns show (left to right): Orthophoto map, physical model bias, bias after ML correction, and bias after ML correction and adding back the mean physical error (*mPE*).

## 4 DISCUSSION

### 4.1 The Promise of Big Data and Machine Learning in Microclimate modelling

With the rise of machine learning in both climate and ecological research, our study demonstrates that microclimate predictions can also benefit from machine learning applications. Microclimate conditions critically influence organismal behaviour and survival (Briscoe et al., 2023; Potter et al., 2013). As microclimate models evolve, they play an increasingly important role in helping ecologists estimate these local conditions accurately (Kearney et al., 2020; Kearney and Porter, 2017; Levy et al., 2016). Reducing prediction biases in these models is essential for enhancing our understanding of how climate variability and change affect biological processes (Levy et al., 2015; Potter et al., 2013; Pincebourde and Woods, 2020). Our study illustrates that machine learning can improve microclimate predictions, as well as highlight parameters that need better representation in physical models, ultimately advancing ecological modelling and decision-making.

Our findings underscore that even a straightforward machine learning algorithm, such as random forests, can provide ecologists and decision-makers with meaningful improvements in microclimate predictions. Such improvements show that ML correction enhances microclimate models by learning complex patterns from observed data, enhancing accuracy beyond what physical models alone can achieve. Already, machine learning is widely applied in large-scale climate models to address bias by identifying complex relationships between model outputs and observed data (Levy and Shahar, 2024,, Table 1A), an approach validated by numerous studies (Chen et al., 2023; Sabarinath et al., 2023; Kesavavarthini et al., 2023; Zhu et al., 2022). For ecological applications, where animals experience conditions at fine spatial scales (Potter et al., 2013; Pincebourde and Woods, 2020; Briscoe et al., 2023; Levy and Shahar, 2024), improving microclimate accuracy is particularly impactful, supporting ecosystem management, agricultural planning, and urban development (Levy and Shahar, 2024).

Integrating ML bias correction with physical modelling opens up significant opportunities. This approach can help refine fine-scale habitat analysis (Suggitt et al., 2011; Pincebourde et al., 2016), reducing biases when identifying areas resilient to temperature fluctuations and crucial for conservation (Anderson et al., 2023; Simonson et al., 2021). Moreover, better microclimate predictions can help anticipate shifts in species distributions (Klinges et al., 2024; Levy and Shahar, 2024; Briscoe et al., 2023; Potter et al., 2013; Pincebourde and Woods, 2020) and animal movements (Sears et al., 2016; Malishev et al., 2018). Such information can make predictions more reliable for early warnings of microclimate changes that may threaten species (Zlotnick et al., 2024; Stark et al., 2023). Together, these advancements provide conservationists and planners with powerful, data-driven tools to better manage the impacts of climate variability on biodiversity.

The growing accessibility of commercial and reliable sensors, coupled with new data analysis methods, presents an exciting opportunity to advance ecological studies and make ecological modelling accessible to a broader range of end-users (Levy and Shahar, 2024; Briscoe et al., 2023). Our current study illustrates this potential: over 33 drone flights were conducted across different seasons, and we collected data covering numerous pixels, which we processed to train our ML model. As drones become more affordable and open-source tools like OpenDroneMap (Gbagir et al., 2023) become increasingly available, implementing our approach in different areas of interest is becoming more practical. Users looking to create high-resolution, ML-corrected microclimate predictions can follow our method by using drone data to map physical structures and vegetation supplemented with meteorological data from local stations or on-line climate resources. To generate initial predictions, these data can be fed into physical microclimate models, such as NicheMapR (Kearney and Porter, 2017) and microclima (Maclean et al., 2019). For model bias correction, users should conduct additional drone flights equipped with thermal cameras to collect empirical temperature data for ML training. For broader-scale applications with coarser spatial resolutions (e.g., tens of meters), satellite data can inform the spatial structure of the environment and provide thermal data for training microclimate models. It is crucial for ecologists and end-users to thoughtfully choose model inputs and features based on their specific research objectives and data collection capabilities, ensuring that their models are both accurate and relevant.

### 4.2 Addressing Biases and Complex Environmental Influences through Machine Learning

Our analysis showed that machine learning can greatly reduce biases in model predictions tied to certain environmental factors. Specifically, the ML model minimized overpredictions at greater heights in open habitats and reduced errors associated with overestimating temperature under high solar radiation and underestimating it under high skyview conditions in shaded areas. While investigating these biases further could help identify potential misspecifications in the physical model, our results suggest that ML can decrease these biases without requiring us to pinpoint the exact underlying assumptions or data inaccuracies responsible for them. For instance, our physical model may overestimate solar radiation in shaded areas, possibly due to an overestimation of diffuse solar radiation (Philipona, 2002), or it may underestimate key processes such as evaporative cooling from nearby vegetation (Lin et al., 2017). Regarding height, we did not account for its effect on wind speed in the physical model, and the fact that wind speed generally increases with height (Campbell and Norman, 1998) may explain some of the observed bias. Similarly, errors related to skyview could arise from an overestimating emissivity or incorrect modelling of longwave radiation (Karsisto and Horttanainen, 2023). It is also possible that the parameters obtained from an online dataset (e.g., albedo, soil temperature, and soil moisture) also contributed to these biases (Zanchi et al., 2023). Identifying these effects can inspire further research to improve physical models or highlight important factors that need accurate measurement for model parameterization. However, for many potential users who lack the resources or expertise to make such refinements, ML offers a practical and effective way to reduce model errors.

Despite the effectiveness of our ML model in reducing many biases, some biases tied to specific environmental factors remained uncorrected. For instance, in open microhabitats, the positive effects of solar radiation and skyview on model biases were not significantly reduced. The reasons for this are unclear, but it may be due to the need for more complex ML models or the insufficiency of our current features to fully capture the complexity of these biases. One missing complexity in our physical model and features is temporal and spatial correlations that may be essential for accurate heat balance predictions. For example, processes like evaporative cooling from nearby vegetation were not included. Additionally, solar radiation and vegetation cover likely exhibit strong temporal dependencies that our spatial model did not consider. For instance, two locations receiving the same solar radiation can have different ground temperatures if one was previously cooler. Furthermore, areas near vegetation experience shifting shade patterns throughout the day; locations that recently transitioned from sun to shade, or vice versa, are particularly susceptible to bias. To better address these complexities, future work could explore models that incorporate spatial relationships, such as ResNet (Liang, 2020), and temporal relationships, such as Recurrent Neural Networks (RNNs) and Long Short-Term Memory networks (LSTMs) (Kaur and Mohta, 2019; Hochreiter and Schmidhuber, 1997). Incorporating features that track shade movement throughout the day or adding information about previous shade conditions may improve model performance. Nonetheless, our current ML model serves as a valuable starting point, offering insights into which features may explain biases and guiding future research on refining microclimate models with more accurate physical processes and parameters.

Another complexity that may cause bias in our shade estimates is the variation in thermal properties among different sources of shade. Accurately modelling microclimates requires distinguishing between shade provided by bushes, rocks, trees, and cliffs, as their thermal characteristics differ significantly. In desert environments, rocks often provide substantially cooler shade temperatures than bushes, and larger rocks are generally cooler than smaller ones (Stark et al., 2023). Modelling these differences requires precise knowledge of each rock and bush’s volume, reflectance, and heat capacity, which all influence the local heat balance (Campbell and Norman, 1998). Additionally, surfaces like tree trunks and cliffs add another layer of complexity, as shade at different heights can exhibit varying temperatures due to higher wind velocities and cooler air temperatures at elevation (Zlotnick et al., 2024; Pincebourde and Woods, 2020). Capturing these variations accurately calls for advanced data collection methods, such as improved drone imaging (e.g., Stark et al., 2022) and the development of drone-based classification techniques (e.g., Ding et al., 2018) that can differentiate between rocks, vegetation, and various shade levels. By incorporating these distinctions, our models could more accurately represent the diverse and complex shading patterns found in ecosystems, significantly enhancing the precision of microclimate predictions.

Our ML model was trained using error values that had been mean-centred, and we observed that adding back the mean error did not significantly improve the model’s predictive performance. While it is possible to calculate the mean error for training purposes when the actual errors are available, this value is not accessible for new, unseen predictions. Therefore, our findings suggest that the model’s ability to reduce bias remains robust, even without incorporating the mean error correction for future predictions. This robustness implies that the model can effectively generalize and minimize biases without needing prior adjustments based on known errors, a desirable characteristic for predictive applications where error distributions may shift over time (Chuang et al., 2020). However, further research could explore whether alternative bias-reduction techniques or adjustments could enhance performance in scenarios with greater variability (Schneider et al., 2023).

### 4.3 Caveats and Future Directions

Although our model demonstrates the potential of machine learning (ML) for improving microclimate estimates, several caveats should be noted. First, not all biases in our microclimate maps were successfully reduced (Fig. 3). Pixels with challenging parameterizations, such as those involving the Triangular Greenness Index (TGI), may have contributed to poor ML corrections. The TGI index, derived from RGB layers, may inaccurately classify dry, less green vegetation as bare ground when using simple thresholding methods (Starý et al., 2020). More accurate parameterization techniques, such as manual annotation, could decrease this bias but would be time-consuming and less feasible for many end-users.

Additional sensors could have improved our remote sensing parameterization. For example, we calculated height as the difference between digital surface model (DSM) and digital terrain model (DTM) maps. However, because the DTM is derived from DSM changes (e.g., Pijl et al., 2020), biases in DTM predictions can reduce height accuracy and thus limit the ML model’s ability to capture microclimate model errors. Employing more direct methods, such as LiDAR measurements—which have been shown to improve terrain accuracy (Oniga et al., 2023; Khanal et al., 2020)—could help reduce these biases in height calculations. Incorporating additional features in the machine learning model that are not currently included in the physical model could also improve ML performance. For example, using spectral data such as the Normalized Difference Vegetation Index (NDVI; reviewed by Huang et al., 2021) has been shown to better capture variations in vegetation cover than RGB-based indices (Furukawa et al., 2021). As remote sensing technologies continue to advance, integrating multiple on-board sensors will likely improve the quality and breadth of environmental data available for future research.

Future efforts should aim to advance our understanding of the ML model’s limitations and strengthen its applicability for predicting microclimates across different environments and temporal scales. One important avenue is collecting additional data within the current habitat to evaluate how biases change across different seasons, providing insights into the model’s temporal robustness. Another step is to gather data from other desert environments to assess the model’s performance across different desert ecosystems and test its ability to generalize within similar, yet distinct, habitats. However, to address undercanopy conditions—particularly in habitats where most of the ground is covered by vegetation, such as forests—our drone-based approach alone may not be sufficient. In these environments, on-the-ground sensors should be deployed beneath different canopy types, species, and heights to capture accurate temperature data. This variation in canopy structure has been shown to significantly affect forest micro-climates (Gril et al., 2023), and can be incorporated into machine learning models to reduce errors in under-canopy microclimate predictions. Expanding data collection to include a range of habitats beyond deserts would also allow for a more robust evaluation of our model’s ability to generalize across diverse ecosystems.

### 4.4 Summary

Machine learning is transforming ecological research by providing powerful predictive tools that can benefit ecologists and decision-makers. We demonstrate the potential of integrating ML with physical models to improve microclimate predictions. Our approach highlights how ML can effectively enhance the accuracy of physical models by correcting biases without requiring extensive investigations into the underlying causes of errors. These advancements offer significant advantages for ecological risk assessments, particularly in the context of climate change and extreme weather events, where accurate microclimate estimates are needed. With the increasing availability of data collection sensors, such as mounted-sensors on drones, and advancements in ML models, we hope ML can be further adapted to improve microclimate modelling. Our findings pave the way for new research directions, emphasizing the value of combining ML and traditional modelling techniques to support more informed conservation and management strategies.

## Declaration of generative AI usage

During the preparation of this manuscript, the authors used ChatGPT to improve language, grammar and clarity. The authors reviewed and edited the suggested corrections and take full responsibility for the content of the published article.

## AUTHOR CONTRIBUTIONS

All authors conceived and designed the study. OL and RDE collected the data. OL led the research and contributed mentorship. MS contributed modelling and validation. AI, IS, RDE, and OL contributed analysis tools. All authors contributed to model development. AI and OL wrote the first draft of the manuscript, and all authors provided com-ments.

## DATA ACCESSIBILITY STATEMENT

Data and code will be made available upon publication.

## ACKNOWLEDGMENTS

We thank Michael Kearney for thoughtful discussions and Simon Jamison for logistical assistance. The research was funded by the National Geographic Society (NGS-84241T-21). There are no conflicts of interest among all authors for this manuscript.

