## Supplementary Information for "From Big Data to Small Scales: Machine Learning Enhances Microclimate Model Predictions"

[2]Tel Aviv University, Center for Artificial Intelligence  Data Science

### Calculating Ground Temperatures Using a Physical Model

At each pixel in the maps, we solved the energy balance equation using Newton-Raphson's iterative method to ensure that the surface fluxes and ground temperatures were in equilibrium. The energy balance equation is given by:

$$S - L - H - E - G = 0, \tag{1}$$

where $S$ represents the solar radiation absorbed by the ground, $L$ denotes the net flux of longwave radiation, $H$ is the flux of sensible heat, $E$ is the flux of latent heat, and $G$ is the flux of soil heat. All fluxes are expressed in units of $W\,m^{-2}$. Positive values for all surface fluxes are directed away from the ground toward the sky, except for solar radiation, which is considered positive when directed toward the ground.

##### Net Solar Radiation

The solar radiation absorbed by the ground ($S$) was calculated as:

$$S = (1 - \alpha)(1 - \text{SHD}) \cdot \text{SWR}, \tag{2}$$

where $\alpha$ represents the albedo in the study area, SHD is a binary indicator of shade conditions (0 for no shade and 1 for shade), and SWR (W $m^{-2}$) is the shortwave radiation reaching the ground. While the albedo is specific to the area, shade conditions and solar radiation were calculated for each 15 $cm^2$ pixel (see details below).

##### Net Longwave Radiation

We calculated the net longwave radiation by adapting the equations from the GitHub documentation of Kearney and Porter (2017) (available at `mrke.github.io/NicheMapR/inst/doc/microclimate-model-theory-equations`). The net flux of longwave radiation ($L$), which is the difference between incoming and outgoing longwave radiation, depends on the longwave downward flux from the sky ($L_{sky}$, W m$^{-2}$), the downward flux from the canopy ($L_{canopy}$, W m$^{-2}$), and the upward flux from the ground ($L_{ground}$, W m$^{-2}$):

$$L = L_{ground} - L_{sky} - L_{canopy} \tag{3}$$

$$L_{ground} = \sigma \epsilon_g T_g^4, \tag{4}$$

$$L_{sky} = SKYV(\sigma \epsilon_{sky} T_a^4 (1 - CLD) + \sigma (T_a - 2)^4) CLD) \tag{5}$$

$$L_{canopy} = \sigma \epsilon_v T_v^4 (1 - SKYV), \tag{6}$$

where $\sigma$ is the Stefan-Boltzmann constant (5.67 $\times$ 10$^{-8}$ W m$^{-2}$ K$^{-4}$), $T_g$ (K) is ground temperature, $T_v$ (K) is vegetation temperature, and $T_a$ (K) is the air temperature. The longwave radiation is also affected by the skyview factor (*SKYV*, i.e., the portion of visible sky, decimal percentage), cloud cover (*CLD*, decimal percentage), and the emissivities of the ground ($\epsilon_g$, decimal percentage) and vegetation ($\epsilon_v$, decimal percentage). For simplicity, we assumed that the vegetation temperature, $T_v$, is the same as the air temperature we measured in the field, measured by our station at a reference height (1.2 m). *CLD* is the cloud cover (decimal percentage), calculated as

$$CLD = \frac{S_g}{S_{g,clean}} \tag{7}$$

where $S_{g,clean}$ is the clear-sky solar radiation, calculated using the *r.sun* function from the GRASS software (GRASS Development Team, 2022). We calculated $\epsilon_{sky}$ as

$$\epsilon_{sky} = 1.72(\frac{e_A}{T_a})^{\frac{1}{7}}, \tag{8}$$

where $e_A$ is the vapor pressure of the air (kPa) (Campbell and Norman, 1998).

**Sensible Heat**

The flux of sensible heat ($H$) was calculated as a function of the coefficient for sensible heat ($c_{sh}$, W m$^{-2}$ K$^{-1}$), ground temperature ($T_g$, K), and air temperatures in the canopy ($T_c$, K), which we assumed equals to $T_a$ at 1.2 m height:

$$H = c_{sh}((T_g - T_c)SHD + (T_g - T_a)(1 - SHD)). \quad (9)$$

As in Levy et al. (2016), we calculated $c_{sh}$ using air density ($\rho$, kg m$^{-3}$), the heat capacity of dry air ($C_p$) at constant pressure (1004.64 J kg$^{-1}$ K$^{-1}$), and the aerodynamic resistance for sensible heat ($r_{ah}$, s m$^{-1}$):

$$c_{sh} = \frac{\rho C_p}{r_{ah}}. \quad (10)$$

The aerodynamic resistance, $r_{ah}$, was derived from Monin-Obukhov similarity theory applied to the surface layer (Brutsaert, 1982; Arya, 1988) and depends on the air temperature at a height of 1.2 m, the wind speed at a height of 10 m, and the roughness height (i.e., the height above the ground at which wind speed is zero). In the model, roughness height was based on the vegetation type.

**Latent Heat Flux**

The flux of latent heat ($E$) was calculated using the coefficient of evaporative heat flux ($c_{ev}$, W m$^{-2}$ Pa$^{-1}$), the saturation vapour pressure at a given ground temperature ($e_s(T_g)$, Pa), relative humidity of the air within soil at the surface ($RH_{soil}$, decimal %, assuming it is equal to the soil moisture, $m^3 m^{-3}$), the vapour pressure of air in the canopy ($e_c$, Pa), and vapour pressure of air above bare ground ($e_g$, Pa):

$$E = c_{ev}((e_s(T_g)RH_{soil} - e_c)SHD + (e_s(T_g)RH_{soil} - e_g)(1 - SHD)). \tag{11}$$

We calculated $c_{ev}$ using air density ($\rho$, kg m$^{-3}$), the aerodynamic resistance for water vapor ($r_{aw}$, s m$^{-1}$), the surface-resistance of ground ($r_s$, s m$^{-1}$), and the psychrometric constant $\gamma$ (Pa K$^{-1}$):

$$c_{ev} = \frac{\rho}{r_{aw} + r_s}. \tag{12}$$

The psychrometric constant was calculated as

$$\gamma = \frac{C_p P_{srf}}{\lambda}, \tag{13}$$

where $P_{srf}$ is the surface pressure (Pa), and $\lambda$ is the latent heat of vaporization or sublimation (J kg$^{-1}$) based on air temperature at 1.2 m above ground, $T_a$: $\lambda$ equalled $h_{vap}$ or $h_{sub}$ when $T_a$ was greater than 0°C or less than 0°C, respectively ($h_{vap} = 2.51 \times 10^6$ and $h_{sub} = 2.84 \times 10^6$). The air density was calculated using the *air.density* function in the *bigleaf* R package (Knauer et al., 2018). For simplicity, as desert plants have evolved adaptations to minimize water loss (Ward, 2016), we assumed that the vapour pressure of air in the canopy equals the vapour pressure of the air above bare ground (i.e., $e_v$ = $e_g$).

**Heat Flux in Soil**

The heat flux to the soil (*G*) was calculated as

$$G = c_{gh}(T_g - T_{soil1}), \tag{14}$$

where $c_{gh}$ is the coefficient of ground heating (W m$^{-2}$ K$^{-1}$), $T_g$ is the ground temperature (K), and $T_{soil1}$ is the soil temperature near the surface at the preceding time (K). We calculated $c_{gh}$ using thermal conductivity ($k$, W m$^{-1}$ K$^{-1}$) of the soil layer based on the temperature and other properties (Niu et al., 2011), and the depth ($\Delta z_1$, m) of the reference layer of soil:

$$c_{gh} = \frac{k}{\Delta z_1}, \tag{15}$$

where $\Delta z_1$ is 6 cm. Soil properties (e.g., porosity and quartz content) were characterized by the characteristics of silty clay soil type in the Noam-MP model (Niu et al., 2011).

### Maps Development for Microclimate Modelling

#### Solar radiation

The solar radiation map represents the total solar radiation (direct and diffuse) that reaches each pixel on the map, measured at a perpendicular angle to the ground surface. To create this map, we used the *r.slope.aspect* command in GRASS GIS to calculate the aspect and slope for each pixel, given the Digital Surface Model (DSM) map, and then applied the *r.sun* command to estimate solar radiation. One of the key inputs for the *r.sun* command is the day of the year and the solar time, which we calculated using the *getSunlightTimes* function from the *suncalc* R package (Thieurmel and Elmarhraoui, 2022). Since *r.sun* calculates solar radiation under clear-sky conditions, we adjusted for cloud cover by multiplying the map by the ratio of the measured solar radiation from the station to the maximum calculated solar radiation on flat surfaces (slope $< 0.1°$).

#### Shade

The shade map is a binary map, with each pixel assigned a value of either 1 (shade) or 0 (no shade). To create this map, we used the *r.sun* command in GRASS GIS to generate an incidence map, which represents the angle of direct sunlight at each pixel. Pixels with NULL values in the incidence map indicate areas in shadow, which we assigned a value of 1 in the shade map.

Importantly, shaded pixels in our analysis do not represent areas directly beneath dense canopy cover (which were excluded from our analysis), but rather areas of bare ground that are temporarily shaded by nearby objects—such as vegetation, rocks, or other landscape features—due to the sun's position at the time of the survey.

#### Skyview

The skyview map represents the skyview factor, which is the proportion of visible sky in each pixel as limited by surrounding physical features such as terrain, rocks, bushes, and trees. We calculated this map using the *r.skyview* command in GRASS GIS. In microclimate modelling, the skyview factor determines the amount of longwave radiation that reaches the ground directly from the sky, with the remainder coming from surrounding physical features. Values range from 0 to 1, with lower values indicating less visible sky (more obstructions) and higher values representing more open areas.

#### Vegetation cover

The vegetation cover map was generated using the Triangular Greenness Index (TGI) for each pixel, which correlates with chlorophyll content (Hunt et al., 2013). Based on our own assessment, we classified pixels with TGI values greater than 0.04 as vegetation, based on visual inspection and calibration against the RGB orthophotos. Although we did not perform a formal sensitivity analysis, this threshold effectively separated vegetated patches from bare ground in our study site. We then excluded these vegetation pixels from the analysis. However, raw TGI values were included as a feature in the machine learning model to improve predictive accuracy. Although vegetation pixels were excluded from the physical model, it may be possible to use canopy temperature models to estimate the temperature of the canopy and, subsequently, the temperature of the ground beneath the canopy.

#### Height

The height map represents the local elevation of surface features relative to the surrounding bare ground for each pixel. To generate this map, we calculated the difference between the Digital Surface Model (DSM), which captures the elevation of all surface features (such as rocks, terraces, and low vegetation), and the Digital Terrain Model (DTM), which represents the underlying bare ground surface. This difference yields a rough approximation of the height for each pixel, reflecting microtopographic variation across the landscape (as suggested in Duffy et al., 2021). For example, in pixels where rocks or vegetation are present, the height value indicates how tall these objects are; in open areas, the value is close to zero.

The height map was not directly used in the physical microclimate model, but height may influence heat-related processes that are not explicitly represented in the physical framework. Therefore, we included the height map as an input feature for the machine learning (ML) model, enabling it to reduce any height-related biases in the physical model that may arise from an oversimplification of the role of height.

### Comparing Online Databases for Parameterizing the Physical Model

To determine which online dataset provides more accurate ground-temperature predictions for our physical model, we compared the model's performance when parameterizing soil temperature, soil moisture, and albedo using either the GLDAS (Rodell et al., 2004) or ERA5 (Hersbach et al., 2020) datasets. We ran the model with each dataset and compared the mean errors (ME) and mean absolute errors (MAE) between the two parameterizations. Our results showed that the GLDAS parameterization yielded more accurate predictions than the ERA5 parameterization (Fig. S3).

#### Comparison Between Our Random Forest and a Deep Learning Model

We fitted a simple deep-learning neural network model (DNN) using a Multi-Layer Perceptron (MLP) architecture. The model included a hidden layer with 10 neurons, a final output layer with 10 neurons, and a maximum of 100 iterations. We selected this model because it serves as a basic neural network framework, is straightforward to manage, and is easy to implement in Python. The DNN model was used to predict the bias of the physical model, analogous to our random forest approach. We used the sklearn python package to run this model, with MLPRegressor library for the DNN implementation. Our results indicated that the random forest model outperformed the neural network model in predicting the biases of the physical model (Fig. S4).

| Symbol | Units | Description | Equation / Source |
|---|---|---|---|
| $\alpha$ | dec. % | Albedo | GLDAS |
| *SHD* | 0 - No<br>1 - Yes | The presence of shade | *r.sun* GRASS command |
| $S_g$ | $Wm^{-2}$ | Solar radiation that reaches the ground | Station |
| *SVF* | dec. % | Skyview | *r.skyview* GRASS command |
| $T_g$ | K | Surface temperature. Initial condition only. | Station |
| $T_a$ | K | Air temperature at 1.2m height | Station |
| *P* | Pascal | Ground pressure | Station |
| *STEMP* | K | Soil temperature in four soil layers: 10, 40, 100, 200 cm | GLDAS |
| *SMOIS* | $m^3\ m^{-3}$ | Soil moisture in four soil layers: 10, 40, 100, 200 cm | GLDAS |
| *RH* | % | Relative humidity | Station |

**Table S1:** Input parameters used in the microclimate models description for physical microclimate model.

| Set | Date | Time |
|---|---|---|
| Train | 18.9.19 | 09:00, 12:00, 14:00, 15:00, 17:20 |
| | 29.5.19 | 08:30, 16:50, 17:30 |
| | 30.1.20 | 09:20, 09:50, 10:50, 12:00, 13:00, 13:50, 14:49, 15:23 |
| | 30.5.19 | 06:00, 06:30 |
| | 7.11.19 | 10:30, 11:00, 13:10, 15:50, 16:40 |
| Test | 12.04.21 | 11:18 |
| | 23.09.19 | 06:10, 07:00, 08:00, 14:10, 15:10, 16:10 |
| | 31.05.21 | 15:16, 17:12, 18:05 |

**Table S2:** The date and time of the different flight surveys. Flights from different dates were selected for training and testing of our bias-correction ML model.

**Table S3:** The effect of machine learning correction on the model's prediction errors (mean, absolute, and squared errors). The table presents the Mean ± SD of all maps before and after applying the machine learning correction. Additionally, it shows the number (N) and percentage of maps where the correction either reduced or increased the mean error. Results are provided for corrections made by the machine learning model both with and without adding back the mean prediction error (mPE).

| | Mean Error | Mean Absolute Error | Mean Squared Error |
|---|---|---|---|
| **Before ML Correction** | 0.560 (±1.993) | 2.750 (±0.849) | 12.129 (±6.612) |
| **After ML Correction** | | | |
| Without mPE | -0.373 (±1.617) | 1.879 (±0.629) | 5.652 (±3.209) |
| With mPE | -0.934 (±1.192) | 1.786 (±0.599) | 5.216 (±2.856) |
| **ML Increased Bias** | | | |
| Without mPE | -1.225 (N=19; 38%) | 2.305 (N=10; 20%) | 8.007 (N=9; 18%) |
| With mPE | -1.295 (N=18; 36%) | 2.415 (N=8; 16%) | 8.106 (N=6; 12%) |
| **ML Reduced Bias** | | | |
| Without mPE | 0.149 (N=31; 62%) | 1.772 (N=40; 80%) | 5.136 (N=41; 82%) |
| With mPE | -0.731 (N=32; 64%) | 1.667 (N=42; 84%) | 4.822 (N=44; 88%) |

**Table S4:** Slopes of the relationships between model features and microclimate bias before (A) and after (B) machine learning correction, and the effect of machine learning correction on the magnitude of these slopes (C). Asterisks represent significant relationships (HDI does not include zero).

| Microhabitat | Feature | Mean ± SD | HDI 2.5% | HDI 97.5% | Significance |
|---|---|---|---|---|---|
| **(A) Physical Model** | | | | | |
| *Open* | TGI | -21.336 ± 2.96 | -26.896 | -15.372 | * |
| | Height | -0.393 ± 0.034 | -0.458 | -0.324 | * |
| | Solar | 0.004 ± 0 | 0.003 | 0.005 | * |
| | Skyview | 2.964 ± 0.396 | 2.17 | 3.718 | * |
| *Shade* | TGI | 4.687 ± 4.373 | -4.105 | 12.998 | |
| | Height | 0.025 ± 0.047 | -0.069 | 0.113 | |
| | Solar | 0.103 ± 0.005 | 0.093 | 0.113 | * |
| | Skyview | -9.645 ± 0.5 | -10.617 | -8.65 | * |
| **(B) After ML Correction** | | | | | |
| *Open* | TGI | 23.495 ± 2.072 | 19.386 | 27.464 | * |
| | Height | -0.1 ± 0.016 | -0.131 | -0.068 | * |
| | Solar | 0.004 ± 0.002 | 0.001 | 0.007 | * |
| | Skyview | 2.686 ± 0.323 | 2.03 | 3.295 | * |
| *Shade* | TGI | 15.245 ± 4.65 | 6.655 | 24.754 | * |
| | Height | 0.039 ± 0.034 | -0.026 | 0.105 | |
| | Solar | 0.056 ± 0.006 | 0.045 | 0.068 | * |
| | Skyview | -5.294 ± 0.6 | -6.496 | -4.164 | * |
| **(C) Effect of ML on Magnitude of Slope** | | | | | |
| *Open* | TGI | -2.159 ± 3.614 | -8.933 | 5.017 | |
| | Height | 0.293 ± 0.038 | 0.218 | 0.366 | * |
| | Solar | 0 ± 0.002 | -0.003 | 0.003 | |
| | Skyview | 0.278 ± 0.511 | -0.694 | 1.29 | |
| *Shade* | TGI | -9.928 ± 5.834 | -20.962 | 1.76 | |
| | Height | -0.001 ± 0.043 | -0.082 | 0.086 | |
| | Solar | 0.047 ± 0.008 | 0.032 | 0.063 | * |
| | Skyview | 4.351 ± 0.777 | 2.792 | 5.808 | * |

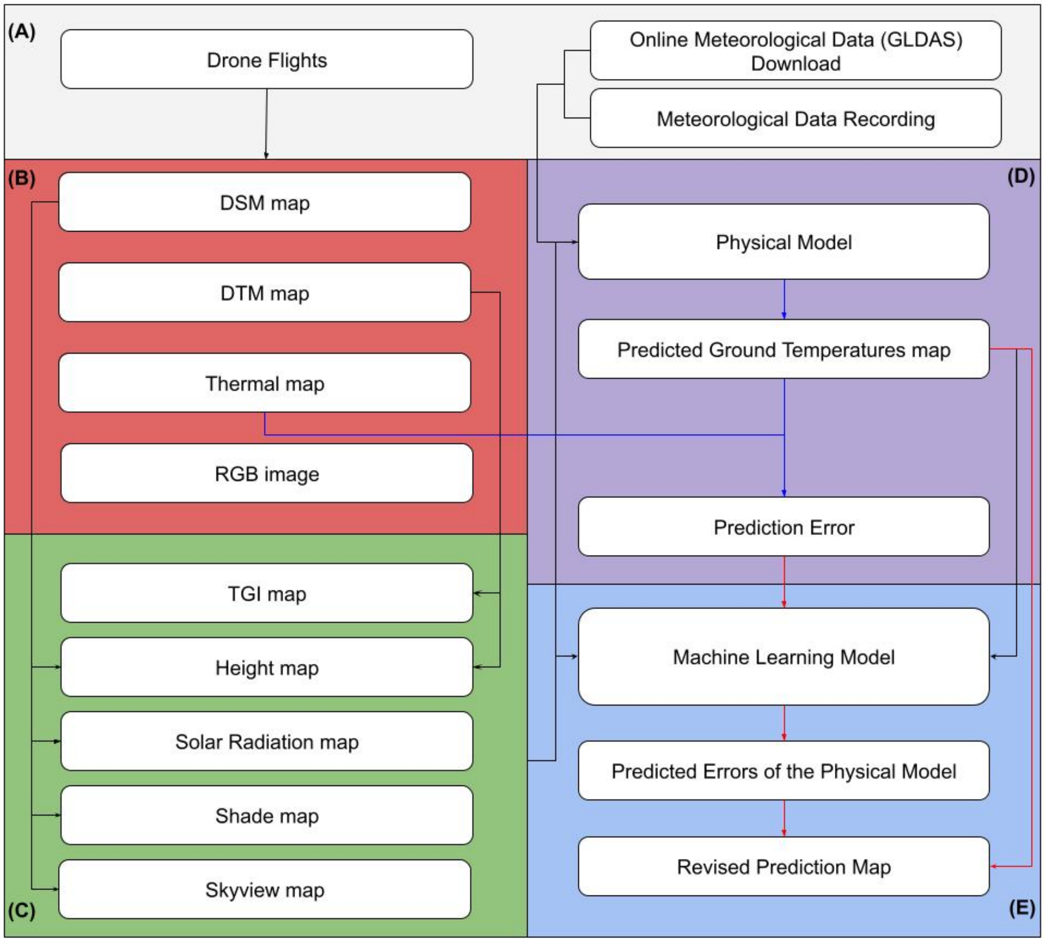


**Figure S1:** Workflow of microclimate modelling. A) Data collection: drone flights and meteorological data (online or recorded). B) Initial image processing: conversion of drone images into Digital Surface Model (DSM), Digital Terrain Model (DTM), orthophotometric (RGB), and thermal maps. C) Feature extraction: generation of TGI, height, real solar, shade, and skyview maps. D) Physical model: input maps and meteorological data are used to produce deterministic prediction maps, and errors are calculated by comparing predictions with thermal maps. E) Machine learning correction: the random forest model predicts the errors of the physical model. These predicted errors are then subtracted from the deterministic predictions to generate revised prediction maps.

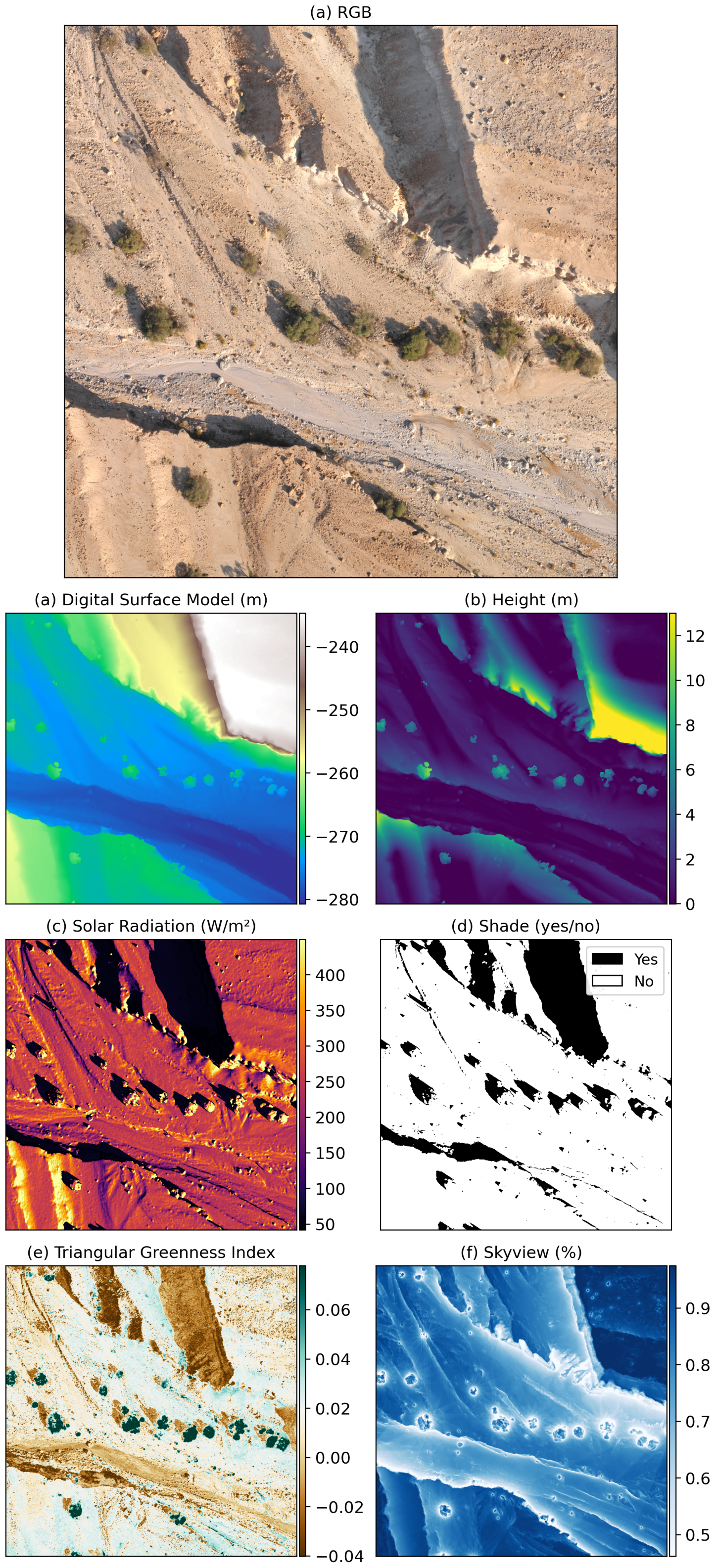


**Figure S2:** Examples of maps used in our study. (a) RGB, (b) Digital Surface Model, (c) height, (d) solar radiation, (e) shade, (f) Triangular Greenness Index, and (g) skyview.

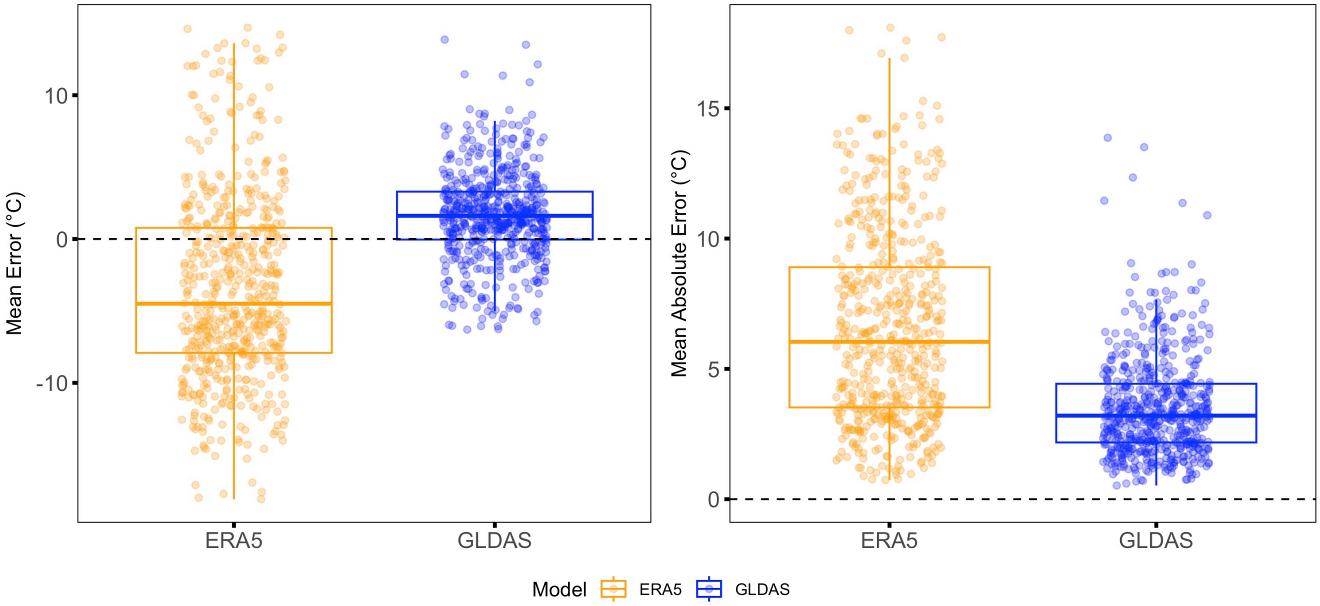


**Figure S3:** Model parameterization using GLDAS produced more accurate predictions of the physical model. The x-axis represents the online database. The left panel represents mean error, and the right panel represents mean absolute error. Each point represents a separate map.

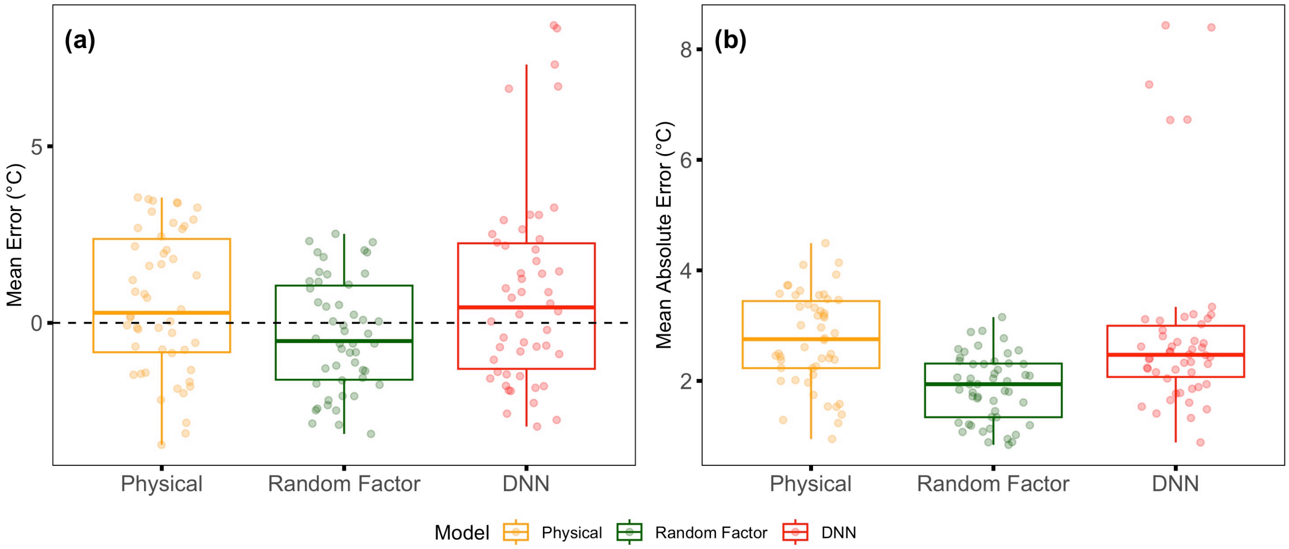


**Figure S4:** The random forest model performed better than a deep neural network (DNN) in correcting the mean errors (a) and mean absolute errors (b). The x-axis represents the model (from left to right: Physical, random forest, and DNN).

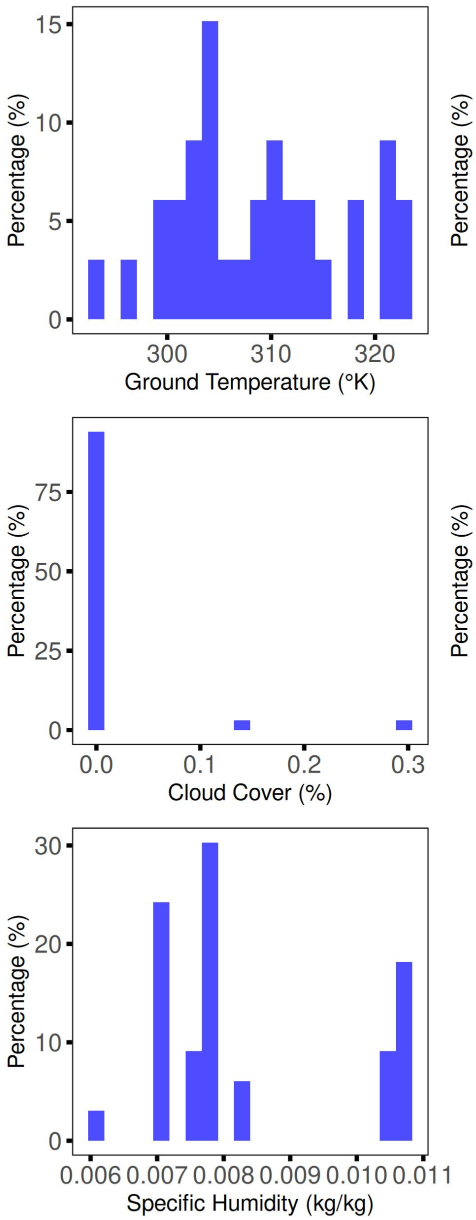


**Figure S5:** The meteorological conditions varied throughout our flight sessions. We show the data distribution of the meteorological variables. Each cell represents a different parameter used in our physical model.

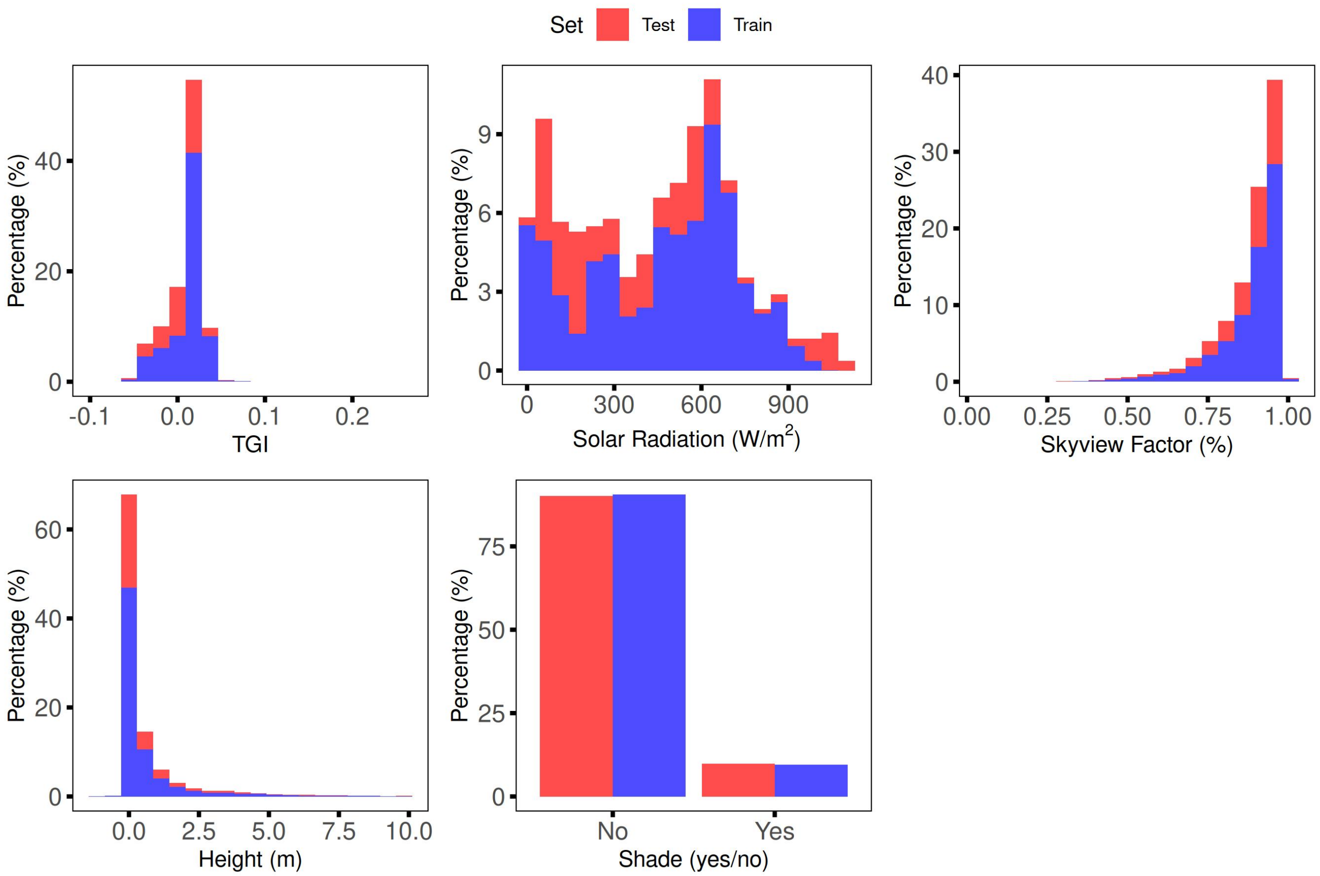


**Figure S6:** Data distribution of model features was similar between train and test maps. Blue and red bars represent train and test sets, respectively.

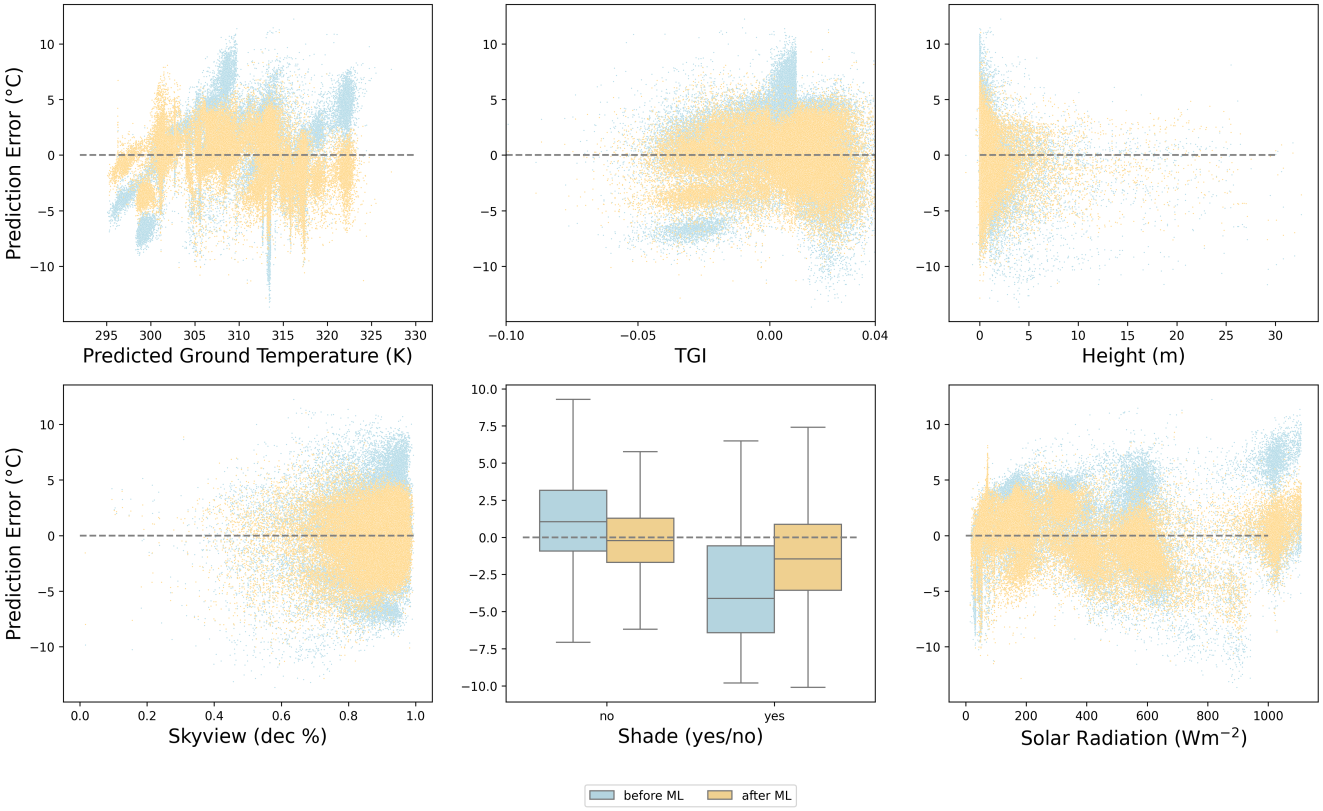


**Figure S7:** Model microclimate bias across each input parameter value. The Y-axis represents the prediction error before (blue) and after (orange) our ML bias correction. X-axes represent the input variable. A) The predictions of the physical model; B) TGI; C) Height; D) Skyview; E) Shade; F) Real solar radiation.

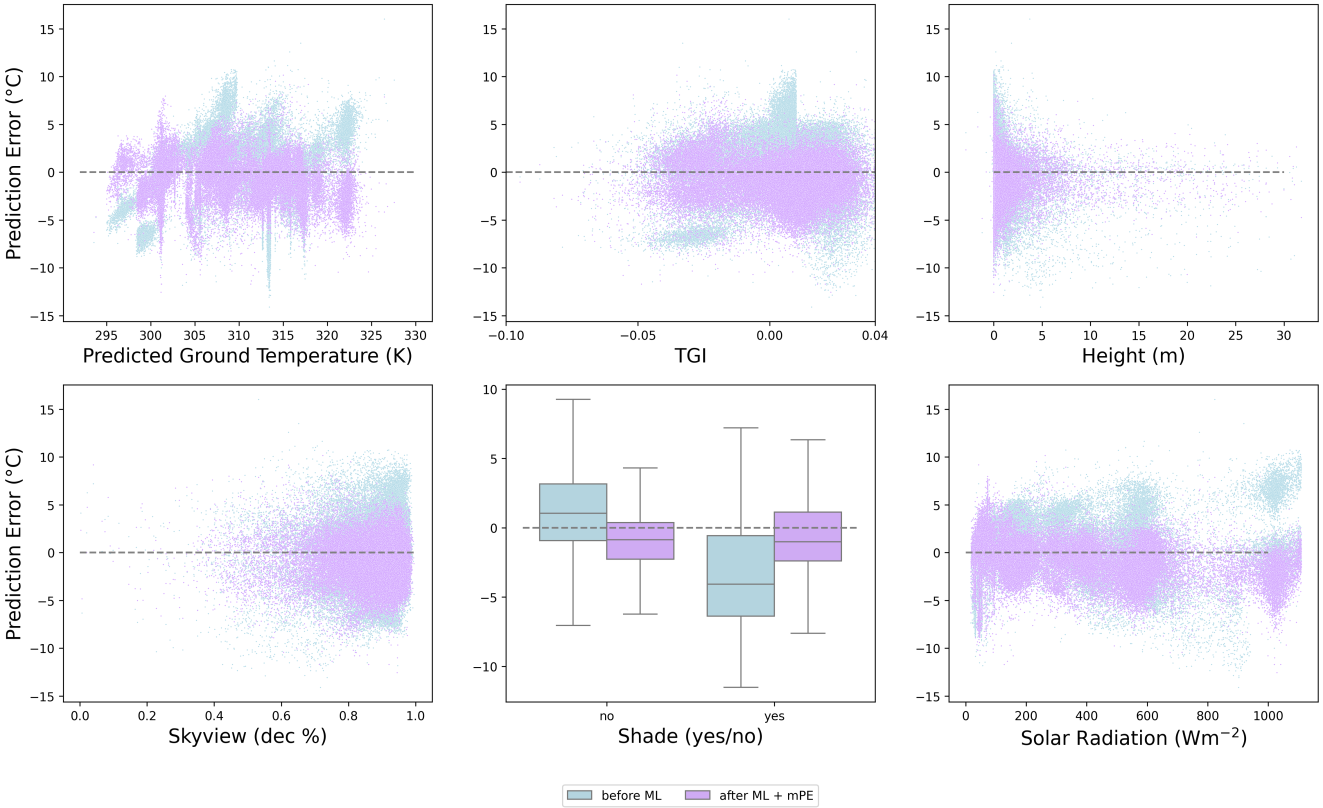


**Figure S8:** Model microclimate bias across each input parameter, before (blue) and after (purple) bias correction after adding back the mean error, mPE. The Y-axis represents prediction error. X-axes represent the input variable. A) Physical prediction; B) TGI; C) Height; D) Skyview; E) Shade; F) Real solar radiation.

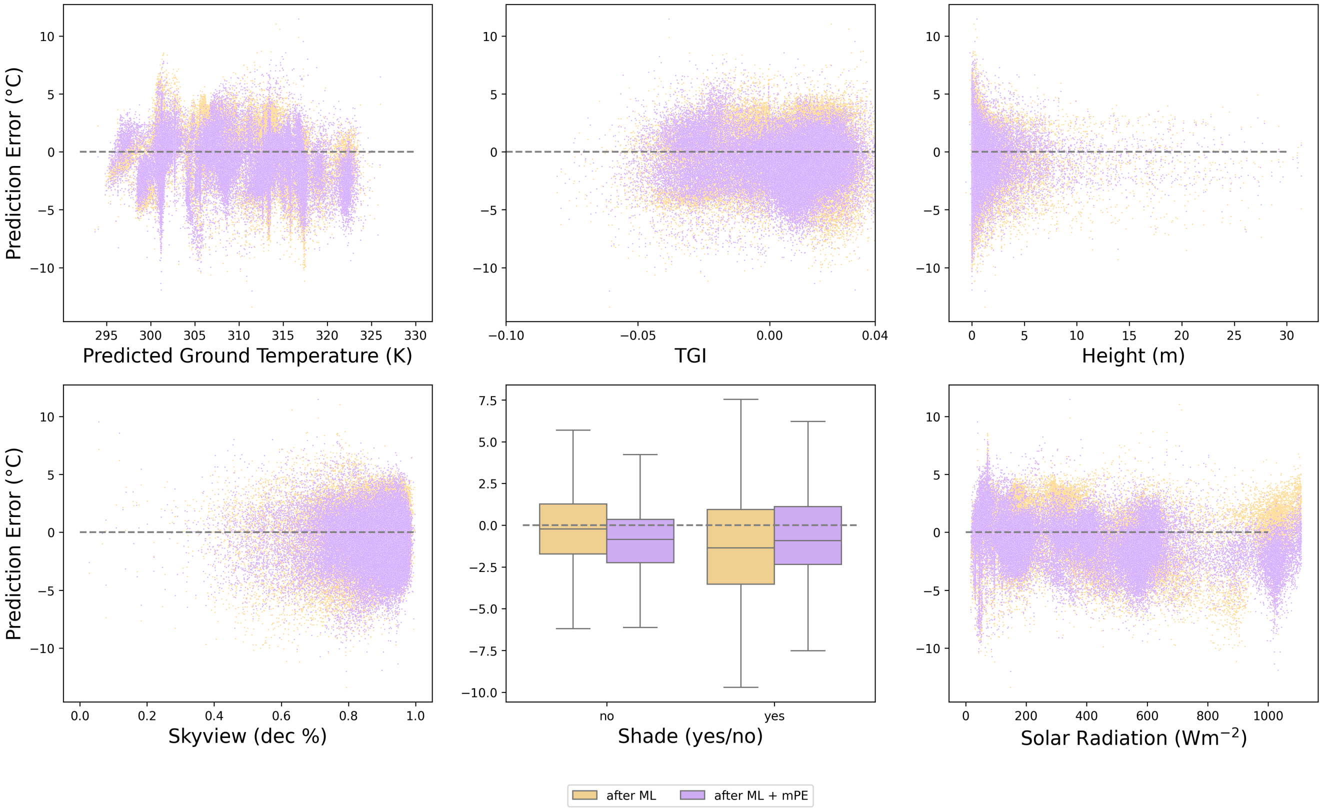


**Figure S9:** Model microclimate bias across each input parameter value after our machine learning, with (purple) and without (orange) adding back the mean error, mPE. The Y-axis represents the prediction error in °C. X-axes represent the input variable. A) Physical prediction; B) TGI; C) Height; D) Skyview; E) Shade; F) Real solar radiation.

Hunt, E. R., P. C. Doraiswamy, J. E. McMurtrey, C. S. Daughtry, E. M. Perry, and B. Akhmedov.

2013. A visible band index for remote sensing leaf chlorophyll content at the canopy scale. International Journal of Applied Earth Observation and Geoinformation, **21**:103–112.

Kearney, M. R. and W. P. Porter. 2017. Nichemapr - an r package for biophysical modelling: the microclimate model. Ecography, **40**:664–674.

Knauer, J., T. S. El-Madany, S. Zaehle, and M. Migliavacca. 2018. Bigleaf - an r package for the calculation of physical and physiological ecosystem properties from eddy covariance data. PLoS ONE, **13**:e0201114.

Levy, O., L. B. Buckley, T. H. Keitt, and M. J. Angilletta. 2016. Dynamically downscaled projections of past and future climates. Ecology, **97**:1888.

Niu, G.-Y., Z.-L. Yang, K. E. Mitchell, F. Chen, M. B. Ek, M. Barlage, A. Kumar, K. Manning, D. Niyogi, E. Rosero, M. Tewari, and Y. Xia. 2011. The community noah land surface model with multiparameterization options (noah-MP): 1. model description and evaluation with local-scale measurements. Journal of Geophysical Research, **116**:D12109.

Rodell, M., P. R. Houser, U. Jambor, J. Gottschalck, K. Mitchell, C.-J. Meng, K. Arsenault, B. Cosgrove, J. Radakovich, M. Bosilovich, J. K. Entin, J. P. Walker, D. Lohmann, and D. Toll. 2004. The global land data assimilation system. Bulletin of the American Meteorological Society, **85**:381 – 394.

Thieurmel, B. and A. Elmarhraoui. 2022. suncalc: Compute Sun Position, Sunlight Phases, Moon Position and Lunar Phase. URL `https://github.com/datastorm-open/suncalc`. R package version 0.5.1.

Ward, D. 2016. The biology of deserts. Oxford University Press.
